# Activin receptor ALK4 coordinates extracellular signals and intrinsic transcriptional programs to regulate development of cortical somatostatin interneurons

**DOI:** 10.1101/807271

**Authors:** Christina Göngrich, Favio A. Krapacher, Hermany Munguba, Diana Fernández-Suárez, Annika Andersson, Jens Hjerling-Leffler, Carlos F. Ibáñez

**Affiliations:** Department of Neuroscience, Karolinska Institute, Stockholm 17177, Sweden; Department of Medical Biochemistry and Biophysics, Karolinska Institute, Stockholm 17177, Sweden; Department of Physiology, National University of Singapore, Singapore 117597, Singapore; Life Sciences Institute, National University of Singapore, Singapore 117456, Singapore; Stellenbosch Institute for Advanced Study, Wallenberg Research Centre at Stellenbosch University, Stellenbosch 7600, South Africa

**Keywords:** GABAergic interneurons, TGF-beta signaling, cerebral cortex, SATB1

## Abstract

Although the role of transcription factors in fate specification of cortical interneurons is well established, how these interact with extracellular signals to regulate interneuron development is poorly understood. Here we show that the activin receptor ALK4 is a key regulator of the specification of somatostatin interneurons. Mice lacking ALK4 in GABAergic neurons of the medial ganglionic eminence (MGE) showed marked deficits in distinct subpopulations of somatostatin interneurons from early postnatal stages of cortical development. Specific loses were observed among distinct subtypes of somatostatin^+^/Reelin^+^ double-positive cells, including Hpse^+^ layer IV cells targeting parvalbumin^+^ interneurons, leading to quantitative alterations in the inhibitory circuitry of this layer. Activin-mediated ALK4 signaling in MGE cells induced interaction of Smad2 with SATB1, a transcription factor critical for somatostatin interneuron development, and promoted SATB1 nuclear translocation and repositioning within the somatostatin gene promoter. These results indicate that intrinsic transcriptional programs interact with extracellular signals present in the environment of MGE cells to regulate cortical interneuron specification.

## Introduction

Network activity in the cerebral cortex is controlled by the interplay of two distinct cell classes: excitatory and inhibitory neurons. While excitatory neurons are long-projecting and use glutamate as their main neurotransmitter, inhibitory neurons are mostly locally-projecting cells (i.e. interneurons) that use gamma amino butyric acid (GABA) to control the activity of excitatory cells (Caputi et al., 2013). In the mouse, cortical GABAergic interneurons are generated during embryonic development in transient neurogenic regions of the basal forebrain known as the medial and caudal ganglionic eminences (MGE and CGE, respectively) and pre-optic area (Hu et al., 2017; Corbin and Butt, 2011). They migrate dorsally following a route tangential to the brain surface to colonize the entire neocortex (Corbin et al., 2001; Marín and Rubenstein, 2003; Wonders and Anderson, 2006). GABAergic interneurons are a diverse, highly heterogenous cell population displaying varied morphological, electrophysiological and molecular characteristics (Markram et al., 2004; Petilla Interneuron Nomenclature Group et al., 2008; Lim et al., 2018). More recently, single-cell gene expression studies have further contributed to our understanding of the molecular diversity of this class of neurons (Tasic et al., 2018; Zeisel et al., 2015). The specification and maturation of the different interneuron subtypes are regulated by the concerted action of genetic and activity-dependent factors (Wamsley and Fishell, 2017). Different combinations of transcription factors function in a sequential fashion to progressively refine the specification of individual interneuron subclasses. MGE-derived somatostatin (SST) expressing interneurons constitute approximately 30% of all cortical GABAergic interneurons and comprise a morphologically (Muñoz et al., 2017), functionally (Hilscher et al., 2017; Xu et al., 2013) and molecularly (Tasic et al., 2018; 2016) diverse class that populates cortical layers II to VI. The development of this subtype is controlled by the sequential activity of the transcription factors Nkx2.1, Lhx6 and SATB1 (Butt et al., 2008; Denaxa et al., 2012; Liodis et al., 2007; Narboux-Nême et al., 2012; Nóbrega-Pereira et al., 2008; Close et al., 2012; Du et al., 2008). While Nkx2.1 and Lhx6 contribute to specify precursors common to both SST and parvalbumin (PV) interneurons, SATB1 is a major determinant of the SST subtype (Close et al., 2012; Denaxa et al., 2012). In agreement with its established role in ventral patterning, Sonic hedgehog (Shh) signaling is required for the early patterning of the basal forebrain as well as Nkx2.1 expression (Fuccillo et al., 2004; Xu et al., 2005). Aside from this early function, it is still not understood how transcriptional programs that specify the distinct types of cortical interneurons interact with extracellular cues present in the environment of MGE cells.

Activin receptor-like kinase 4 (ALK4) is a type I serine-threonine kinase receptor for a subset of TGFβ superfamily ligands, that includes activins A, and B, as well as members of the Growth Differentiation Factor (GDF) subfamily, including GDF-1, −3 and −5, among others (Schmierer and Hill, 2007; Moustakas and Heldin, 2009; Dijke et al., 1994). Ligand binding to type II receptors ActrIIA or ActrIIB recruits ALK4 to the complex, resulting in ALK4 GS-domain phosphorylation and activation of the ALK4 kinase (Budi et al., 2017; Massagué, 2012). Similar to other type I receptors, canonical signaling by ALK4 involves the recruitment and phosphorylation of Smad proteins 2 and 3, which then partner with Smad 4 and translocate to the cell nucleus, where they regulate gene expression in conjunction with cell type specific transcription factors (Schmierer and Hill, 2007; Budi et al., 2017; Massagué, 2012). ALK4 is expressed in the extra-embryonic ectoderm and in the epiblast of early post-implantation embryos; by midgestation, its expression is nearly ubiquitous, including the central nervous system (Gu et al., 1998; Verschueren et al., 1995). A null mutation in mouse *Acvr1b*, the gene encoding ALK4, is embryonic lethal due to defects in primitive streak formation and gastrulation (Gu et al., 1998), and no conditional approaches have yet been reported to address the function of this receptor in nervous system development or function. Despite the importance of the ALK4 signaling system in the development of many peripheral organs and tissues, there is a large gap of knowledge on its possible functions in the brain.

In this study, using different lines of conditional mutant mice, we show that ALK4 is indispensable for the development of cortical SST-expressing GABAergic interneurons. ALK4 signaling is required before the acquisition of the SST phenotype and contributes to the nuclear localization and function of SATB1, a critical transcriptional determinant of SST interneuron specification. A marked paucity of SST interneurons in layer IV resulted in abnormal activity of the inhibitory circuit in this cortical layer. These results define a previously unknown function of ALK4 in the basal forebrain, linking intracellular transcription factor cascades known to govern interneuron development to the extracellular environment of the developing brain.

## Results

### Loss of SST^+^ cortical interneurons after deletion of ALK4 in GABAergic cells of the MGE

In order to dissect cell-autonomous functions of ALK4 in specific tissues, we generated a conditional null allele of the mouse *Acvr1b* gene (*Alk4*^fl^) with loxP sites flanking exons 5 and 6, encoding essential regions of the ALK4 kinase domain (Figure S1). Gene deletion in GABAergic neurons was achieved by crossing *Alk4*^fl/fl^ and *Gad67*^Cre^ mice. Earlier radioactive *in situ* hybridization studies had shown low but widespread expression of *Alk4* mRNA in the developing basal forebrain (Verschueren et al., 1995). In our hands, conventional histological methods with standard riboprobes or commercially available antibodies did not give adequate results. However, using RNAscope *in situ* hybridization, a method of much greater sensitivity, we succeeded to detect *Alk4* mRNA expression in the mantle zone of the MGE in wild type E12.5 mouse embryos (Figure 1). The signal for *Alk4* mRNA overlapped with that of *Lhx6* mRNA (Figure 1), a specific marker of MGE-derived, postmitotic GABAergic interneurons (Liodis et al., 2007; Du et al., 2008). *Alk4* mRNA levels in the ventricular and subventricular zones, containing proliferating precursors labeled by PCNA staining, were very low (Figure 1), indicating a predominantly postmitotic expression of *Alk4* mRNA in the mouse MGE. Importantly, the *Alk4* mRNA signal was significantly reduced in the MGE mantle zone of *Gad67*^Cre^:*Alk4*^fl/fl^ mutant embryos, (Figure S2). We also detected significant levels of mRNAs encoding various ALK4 ligands, including *InhbA* (for activin A), *InhbB* (for activin B), and *Gdf1* (for Growth and Differentiation Factor 1), in the subpallium of E12.5 and E14.5 *Alk4* mutant and control embryos (Figures S3A to C).

**Figure 1.**
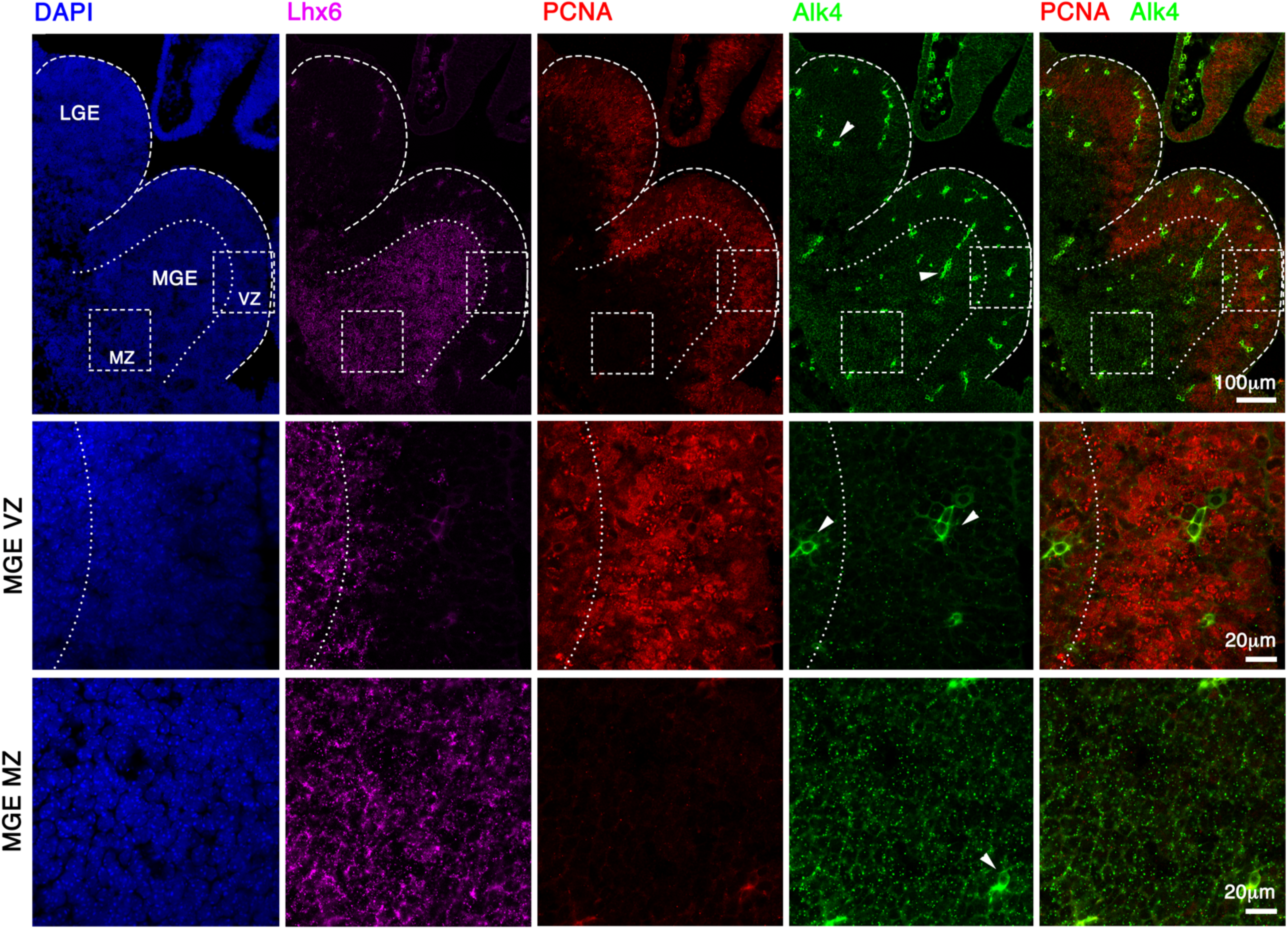
*Alk4* mRNA expression in the mantle zone of the MGE. RNAscope *in situ* hybridization analysis of *Alk4* mRNA expression (green) in the ventricular/subventricular (VZ) and mantle (MZ) zones of the MGE in wild type E12.5 mouse embryos. Upper row shows lower magnification images. Middle and lower rows show higher magnification images of ventricular (VZ) and mantle (MZ) zones, respectively, of areas boxed in upper panel. *In situ* hybridization for *Lhx6* mRNA was used as marker for postmitotic interneurons of the MGE (purple). Immunohistochemistry for PCNA (red) was used to mark proliferating cells in the VZ. COunterstaiing with DAPI is shown in blue. Note that the *Alk4* mRNA signal overlaps with the area labelled by *Lhx6* mRNA in the mantle zone, but not with that labeled by PCNA in the proliferative zone. Arrowheads point to blood vessels, which show unspecific signal in the green channel. Scale bars: 100µm (upper row); 20µm (middle and lower rows).

We used a *Rosa26*^tdTom^ reporter allele to assess recombination efficiency driven by *Gad67*^Cre^ among cortical GABAergic interneurons. We found that 80 to 90% of cells expressing either SST, PV, Reelin (RELN) or vasoactive intestinal peptide (VIP) were positive for tdTomato in the cortex of postnatal day 30 (P30) *Gad67*^Cre^:*Rosa26*^tdTom^ mice (Figures S4A and B), indicating high levels of recombination. Loss of ALK4 in GABAergic cells (*Gad67*^Cre^:*Rosa26*^tdTom^:*Alk4*^fl/fl^ mice) resulted in 34.6 ± 4% reduction in tdTomato^+^ cells in the P30 somatosensory cortex compared to control mice (*Gad67*^Cre^:*Rosa26*^tdTom^:*Alk4*^+/+^), with cell losses in all layers, except layer I (Figures 2A and B). We assessed cell numbers among different classes of cortical interneurons, including non-overlapping subpopulations of MGE-derived SST^+^ and PV^+^ neurons, VIP^+^ neurons derived from the caudal ganglionic eminence (CGE), and RELN^+^ neurons derived from both the MGE and CGE (Figure 2C to G). In these experiments, we compared mutant *Gad67*^Cre^:*Alk4*^fl/fl^ mice to three types of controls, namely wild type, *Gad67*^Cre^:*Alk4*^+/+^ and *Alk4*^fl/fl^ mice. A small reduction in PV^+^ cells was detected in cortical layer IV of the mutants when compared to wild type mice, but this difference was not statistically significant when compared to the two other control groups (Figures 2C and D). On the other hand, SST^+^ cell numbers were strongly reduced (between 60 and 70%) in all cortical layers of the mutants compared to all three control lines (Figure 2E). RELN^+^ cells were also reduced (Figure 2F), while no major differences were observed in VIP^+^ cells, except for a small increase in layer V, when compared to wild type and *Gad67*^Cre^ but not *Alk4*^fl/fl^ mice (Figure 2G). No differences were observed in the number of SST^+^ neurons in the striatum of P30 mutant mice (Figure S5).

**Figure 2.**
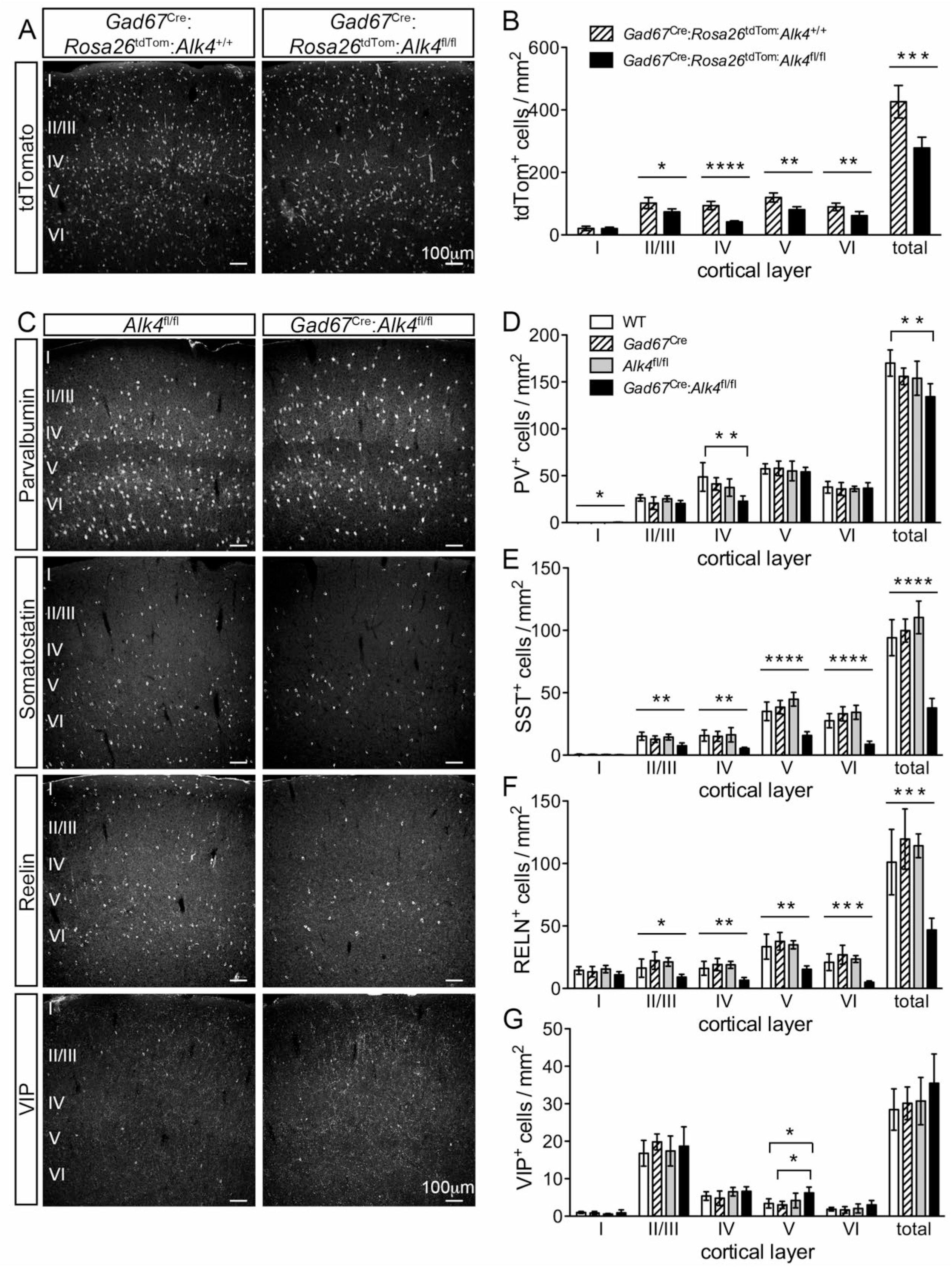
Loss of SST^+^ cortical interneurons after deletion of ALK4 in GABAergic cells of the MGE. (A) Representative confocal images of tdTomato fluorescence in the somatosensory cortex of P30 *Gad67*^Cre^:*Rosa26*^tdTom^:*Alk4*^+/+^ mice (left) and *Gad67*^Cre^:*Rosa26*^tdTom^:*Alk4*^fl/fl^ mice (right). Roman numerals indicate cortical layers as detected by DAPI nuclear counterstaining (not shown here). Scale bars: 100µm. (B) Quantification of tdTomato^+^ cells in the P30 somatosensory cortex of the indicated genotypes. Cell counts were normalized to the area of the section that was imaged and averaged per mouse. Data is presented as mean ± SD. N = 5 mice per genotype. *, P ≤ 0.05; **, P ≤ 0.01; ***, P ≤ 0.001; ****, P ≤ 0.0001 (Student’s t-test). (C) Representative confocal images of the somatosensory cortex of P30 control (*Alk4*^fl/fl^, left) and mutant (*Gad67*^Cre^:*Alk4*^fl/fl^, right) mice showing fluorescence immunohistochemical detection of PV, SST, RELN and VIP expression. Roman numerals indicate cortical layers as detected by DAPI nuclear counterstaining (not shown here). Scale bars: 100 µm. (D to G) Quantification of PV^+^ (D), SST^+^ (E), RELN^+^ (F) and VIP^+^ (G) cells in the P30 somatosensory cortex of the indicated genotypes. Cell counts were normalized to the area of the section that was imaged and averaged per mouse. Data is presented as mean ± SD. N = 5 mice per genotype. P values determined by ANOVA (Bonferroni post-hoc test) are indicated as in panel (B). Statistically significant differences between *Gad67*^Cre^:*Alk4*^fl/fl^ mice and all three control grpups are indicated in (E), (F) and (G). In panel (D), statistically significant differences were only observed between *Gad67*^Cre^:*Alk4*^fl/fl^ mice and the wild type group.

With the exception of layers II/III, comparable losses of SST^+^ cells were observed in the P30 cortex of *Nkx2.1*^Cre^:*Alk4*^fl/fl^ mice (Figures 3A and B), which specifically targets proliferating precursors of GABAergic cells residing in the SVZ of the MGE (Xu et al., 2008). As indicated by earlier studies (Xu et al., 2008), *Nkx2.1*^Cre^ recombines in only 60% of SST^+^ cells in layers II-IV of the neocortex, and it is therefore likely that a large part of the SST^+^ interneurons that were lost after *Alk4* deletion using the *Gad67*^Cre^ line were spared in the *Nkx2.1*^Cre^ line. Interestingly, we observed no cell loses in *Sst*^IRES*-* Cre^:*Alk4*^fl/fl^ mice (Figures 3C and D), which induces recombination at later stages and exclusively in SST-expressing cells. In our hands, only a small proportion of MGE-derived interneurons expressed Cre from the *Sst*^IRES*-*Cre^ locus 48h after their generation, while the vast majority expressed Cre in the *Gad67*^Cre^ mouse within this period (Figures S6A and B), in accordance with the later onset of *Sst*^IRES*-*Cre^ locus activation. Together, these results suggest that ALK4 is required in MGE-derived SST interneuron precursors that have just left the cell cycle but not yet begun expression of SST. Importantly, the loss of tdTomato^+^ cells in *Gad67*^Cre^:*Rosa26*^tdTom^:*Alk4*^fl/fl^ mice suggests a loss of interneurons in the mutant cortex, rather than downregulation of marker gene expression. The normal numbers of SST^+^ cells in the P30 cortex of *Sst*^IRES*-*Cre^:*Alk4*^fl/fl^ mice also indicated that ALK4 is dispensable for the maintenance of the SST phenotype.

**Figure 3.**
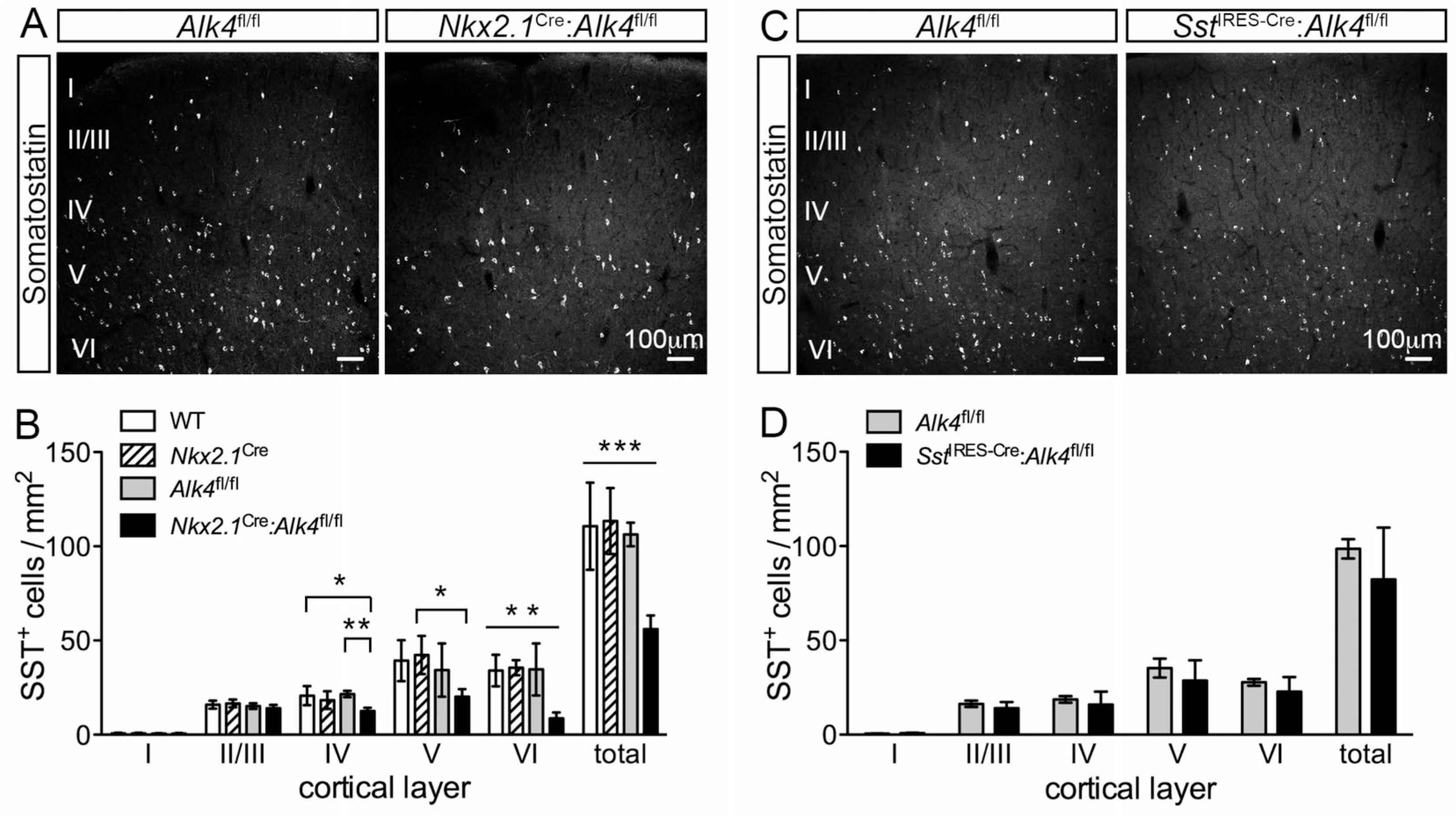
Loss of SST^+^ cortical interneurons after ALK4 deletion with *Nkx2.1*^Cre^but not *Sst*^IRES*-*Cre^. (A) Representative confocal images of fluorescence immunohistochemical detection of SST^+^ cells in the somatosensory cortex of P30 control *Alk4*^fl/fl^ (left) and mutant *Nkx2.1*^Cre^:*Alk4*^fl/fl^ (right) mice. Roman numerals indicate cortical layers as detected by DAPI nuclear counterstaining (not shown here). Scale bars: 100 µm. (B) Quantification of SST^+^ cells in the P30 somatosensory cortex of the indicated genotypes. Cell counts were normalized to the area of the section that was imaged and averaged per mouse. Data is presented as mean ± SD. N = 5 mice per genotype. *, P ≤ 0.05; **, P ≤ 0.01; ***, P ≤ 0.001; ****, P ≤ 0.0001 (ANOVA, Bonferroni post-hoc test). (C) Representative confocal images of fluorescence immunohistochemical detection of SST^+^ cells in the somatosensory cortex of P30 control *Alk4*^fl/fl^ (left) and *Sst*^IRES-Cre^:*Alk4*^fl/fl^ (right) mice. Roman numerals indicate cortical layers as detected by DAPI nuclear counterstaining (not shown here). Scale bar: 100 µm. (D) Quantification of SST^+^ cells in somatosensory cortex of *Alk4*^fl/fl^ and *Sst*^IRES-Cre^:*Alk4*^fl/fl^ mice. Data is presented as mean ± SD. N = 5 mice per genotype. There were no statistically significant differences (P > 0.05, Student’s t-test).

### Distinct subpopulations of SST^+^ cortical interneurons in layers IV, V and VI are affected by ALK4 loss in GABAergic cells of the MGE

Previous studies have indicated some overlap between SST and RELN expression in subpopulations of MGE-derived GABAergic interneurons (Pesold et al., 1999; Miyoshi et al., 2010). In our hands, approximately two thirds of all SST^+^ interneurons in the somatosensory cortex of the P30 mouse brain co-expressed RELN (Figures 4A to C). This was also the subpopulation mostly affected in *Gad67*^Cre^:*Alk4*^fl/fl^ mice, showing significant losses of double-positive neurons in cortical layers II to VI (Figure 4B). In contrast, SST^+^/RELN^-^ cells were not as severely affected in these mutants, with only a relatively smaller reduction in layer VI (Figure 4C). In addition, all cortical layers showed reduced numbers of RELN^+^ neurons that lacked SST expression (Figure 4D). As these cells are thought to be derived from the CGE (Miyoshi et al., 2010), we performed a similar analysis in *Nkx2.1*^Cre^:*Alk4*^fl/fl^ mice, which, as mentioned above, specifically targets GABAergic precursors derived from the MGE (Figures 4E to G). There were no significant loses of SST ^-^/RELN^+^ cells in the cortex of these mice, except for a small group of cells in layer V (Figure 4G), suggesting non-cell-autonomous effects of ALK4 on this subpopulation. The loss of SST^+^ interneurons in *Nkx2.1*^Cre^:*Alk4*^fl/fl^ mice was comparable to that observed in *Gad67*^Cre^:*Alk4*^fl/fl^ mice (Figures 4E and F), in agreement with the MGE origin of these cells.

**Figure 4.**
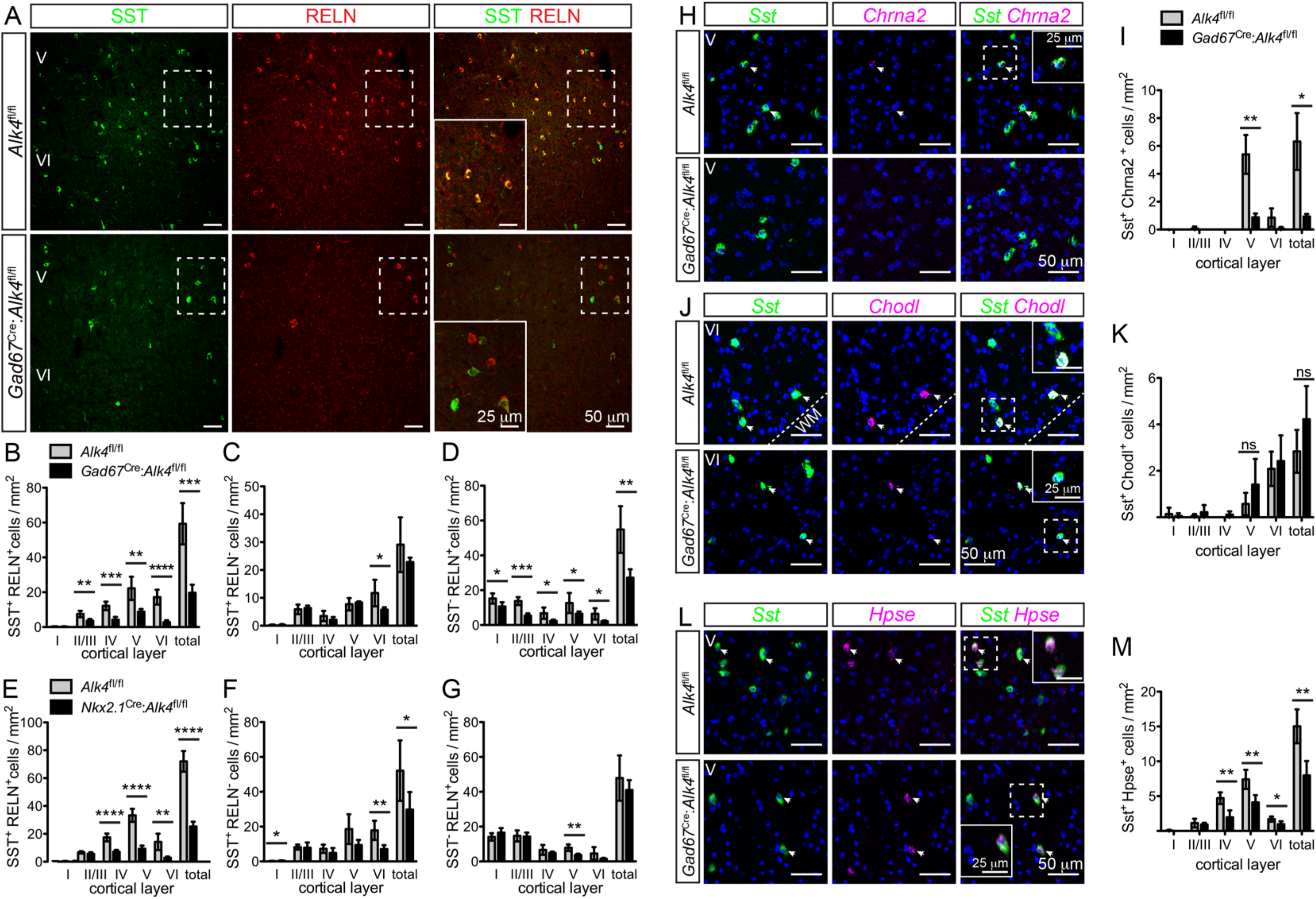
Distinct subpopulations of SST^+^ cortical interneurons are affected by ALK4 loss in GABAergic cells of the MGE. (A) Representative confocal images of fluorescence immunohistochemical detection of SST (green) and RELN (red) in the somatosensory cortex of P30 control *Alk4*^fl/fl^ (top row) and mutant *Gad67*^Cre^:*Alk4*^fl/fl^ (bottom row) mice. Insets show magnified views of the areas outlined by the dashed boxes. Roman numerals indicate cortical layers as detected by DAPI nuclear counterstaining (not shown here). Scale bars: 50 µm (main images), 25 µm (insets). (B-D) Quantification of SST/RELN double positive (B), SST^+^ RELN^-^ (C) and SST^-^ RELN^+^ (D) cells in control *Alk4*^fl/fl^ (grey bars) and mutant *Gad67*^Cre^:*Alk4*^fl/fl^ (black bars) mice. Cells were counted in confocal images spanning layers I to VI of the P30 somatosensory cortex. Cell counts were normalized to the area of the section that was imaged and averaged per mouse. Data is presented as mean ± SD. N = 5 mice per genotype. *, P ≤ 0.05; **, P ≤ 0.01; ***, P ≤ 0.001; ****, P ≤ 0.0001 (Student’s t-test). (E-G) Similar analysis as (B-D) performed in *Alk4*^fl/fl^ (grey bars) and mutant *Nkx2.1*^Cre^:*Alk4*^fl/fl^ (black bars) mice. (H) Representative confocal images of RNAScope *in situ* hybridization for *Sst* (green) and *Chrna2* (red) mRNAs in layer V of the somatosensory cortex of P30 control *Gad67*^Cre^:*Alk4*^fl/+^ (top row) and mutant *Gad67*^Cre^:*Alk4*^fl/fl^ (bottom row) mice. Inset show magnified view of the area outlined by the dashed box. DAPI nuclear counterstaining is shown in blue. Arrowheads indicate double positive cells. Scale bars, 50µm (main images), 25µm (insets).. (I) Quantification of double positive cells for *Sst* and *Chrna2* mRNAs in cortical layer V of control *Gad67*^Cre^:*Alk4*^fl/+^ (grey bars) and mutant *Gad67*^Cre^:*Alk4*^fl/fl^ (black bars) mice. Cell counts were normalized to the area of the section that was imaged and averaged per mouse. Data is presented as mean ± SD. N = 5 mice per genotype. *, P ≤ 0.05; **, P ≤ 0.01 (Student’s t-test). (J, K) Similar analysis as (H, I) on double positive cells for *Sst* and *Chodl* mRNAs. VI, cortical layer VI; WM, white matter. ns, not statistically significant difference (Student’s t-test). (L, M) Similar analysis as (H, I) on double positive cells for *Sst* and *Hpse* mRNAs. V, cortical layer V. *, P ≤ 0.05;; **, P ≤ 0.01 (Student’s t-test).

Although highly significant, the loss of SST^+^ cortical interneurons in *Alk4* mutant mice was not complete. This could have been due to a stochastic requirement for ALK4 signaling across different SST^+^ interneuron subtypes, or else reflect the existence of subpopulations of SST^+^ cells with distinct requirements. Recent studies have leveraged single-cell RNA-Seq methods to molecularly define distinct populations of cortical GABAergic interneurons, including several subtypes of SST^+^ cells (Tasic et al., 2018; Mayer et al., 2018; Mi et al., 2018; Naka et al., 2018). Although much remains to be learnt about the functional features of many of those subpopulations, a few of the molecular markers identified do label cortical GABAergic neurons with known functional properties. Expression of *Chrna2* (encoding Cholinergic Receptor Nicotinic Alpha 2 Subunit) labels a subpopulation of classical SST^+^ Martinotti cells in layer V (Tasic et al., 2018; 2016) that extend axons to projection neurons in layer I (Hilscher et al., 2017). Chondrolectin (encoded by *Chodl*) has been found to mark a distinct class of deep layer SST^+^ cells (Tasic et al., 2018; 2016) with long-range projections that cross to the contralateral hemisphere (Taniguchi et al., 2011; Kubota et al., 1994; Tomioka et al., 2005). Finally, *Hpse* mRNA (encoding Heparanase) was also recently found to specifically label a subpopulation of SST^+^ cells in cortical layer IV that targets PV^+^ fast-spiking interneurons in the same layer (Naka et al., 2018). This subpopulation is sometimes referred to as X94 cells, since a large fraction of these cells is labeled in the X94 *Gad67*^eGFP^ mouse transgenic line (Ma et al., 2006). Using RNAscope *in situ* hybridization, we investigated expression of *Chrna2*, *Chodl* and *Hpse* mRNAs in the cerebral cortex of P30 *Gad67*^Cre^:*Alk4*^fl/fl^ mutant mice and *Gad67*^Cre^:*Alk4*^fl/+^ heterozygote controls (Figures 4H to M). We found that the SST^+^ subpopulation of Martinotti cells expressing *Chrna2* mRNA was almost entirely ablated in the mutants (Figures 4H and I). On the other hand, long-projecting SST^+^ neurons expressing *Chodl* mRNA were spared (Figures 4J and K). Finally, a significant proportion of SST^+^ cells expressing *Hpse* mRNA, up to 50% in layer IV, was also lost in the *Alk4* homozygote mutants (Figures 4L and M). Together, these results suggest a selective dependence on ALK4 signaling among distinct subtypes of SST^+^ interneurons for their correct development.

### Altered inhibitory circuitry in cortical layer IV of mice lacking ALK4 in GABAergic cells of the MGE

Electrophysiolgical and morphological characterization of layer IV SST^+^ interneurons has shown that, rather than targeting pyramidal neurons, many of these cells contact layer IV PV^+^ interneurons (Ma et al., 2006; Muñoz et al., 2017; Xu et al., 2013), thereby gating the inhibitory action of PV^+^ cells on pyramidal neurons. As shown above, a significant fraction of these cells, expressing *Hpse* mRNA, was lost in the *Gad67*^Cre^:*Alk4*^fl/fl^ mutants (Figures 4L and M). Also *Nkx2.1*^Cre^:*Alk4*^fl/fl^ mice lacked a sizable fraction of SST^+^ cells in this layer (Figure 4E). We therefore sought to assess whether inhibition of layer IV PV^+^ cells was affected in these mice. To this end, we performed whole cell patch clamp recordings in layer IV PV^+^ cells of P30 *Nkx2.1*^Cre^:*Rosa26*^tdTom^ mice. Layer IV PV^+^ cells were identified by tdTomato expression, as well as their characteristic soma shape and spiking pattern (Figures 5A and B). We compared recordings made in homozygote *Alk4* mutant mice (*Nkx2.1*^Cre^:*Alk4*^fl/fl^:*Rosa26*^tdTom^) to equivalent measurements obtained in heterozygote control mice (*Nkx2.1*^Cre^:*Alk4*^fl/+^:*Rosa26*^tdTom^). We found that both the amplitude and frequency of inhibitory post-synaptic currents (IPSCs) in layer IV PV^+^ neurons were greatly reduced (Figures 5C and D), suggesting that a large proportion of the inhibitory input to these cells may be provided by SST^+^ layer IV interneurons that depend on ALK4 signaling for their development.

**Figure 5.**
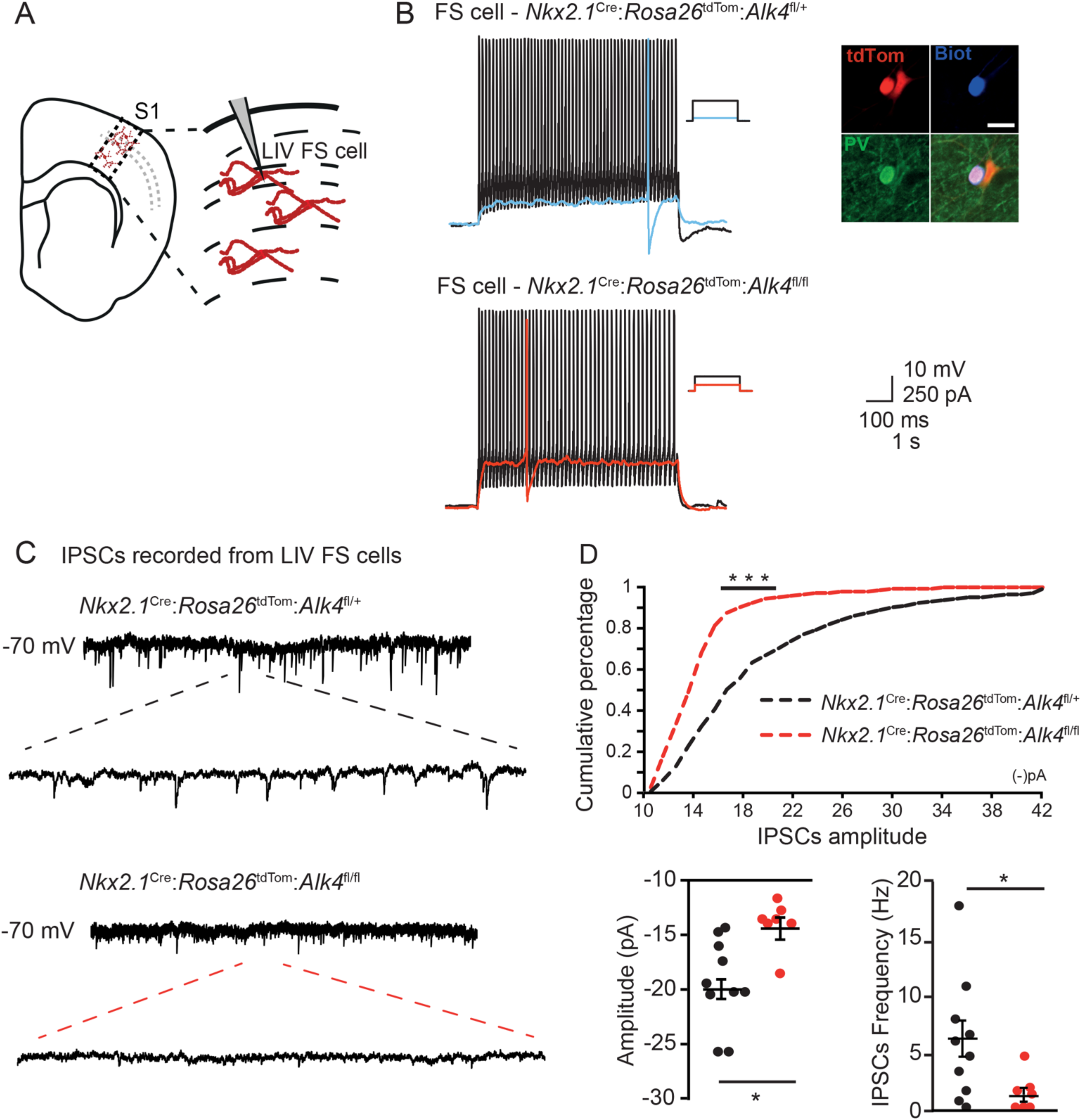
Altered inhibitory circuitry in cortical layer IV of mice lacking ALK4 in GABAergic cells of the MGE. (A) Schematic representation of patch-clamp experimental setting. (B) Representative traces of layer IV fast-spiking cells in heterozygote control (*Nkx2.1*^Cre^:*Alk4*^fl/+^:*Rosa26*^tdTom^) and homozygote mutant (*Nkx2.1*^Cre^:*Alk4*^fl/fl^:*Rosa26*^tdTom^) mice. Top right panel shows post-hoc fluorescence detection of tdTomato (red), PV (green) and neurobiotin (Biot, blue) confirming their co-localization. (C) Fast-spiking (FS) cell voltage-clamp traces recorded under glutamatergic blockade in order to isolate inhibitory postsynaptic currents (IPSCs). (D) Cumulative distribution of IPSCs’ amplitudes in FS-cells of control (black) and *Alk4* mutant (red) mice (top). ***, P≤0.001, Unpaired Kologorov-Smirnov test for continuous probability distributions. Below panels show amplitude (left) and frequency (right) of FS-cells IPSCs in control (black) and *Alk4* mutant (red) mice. Error bars show mean ± SEM. N=10; *, P ≤ 0.05; (Student’s two-tailed t-test).

### Perinatal loss of SATB1^+^ interneuron precursors in *Gad67*^Cre^:*Alk4*^fl/fl^ mutant mice

Comparison of the phenotypes induced by *Gad67*^Cre^, *Nkx2.1*^Cre^ and *Sst^IRES-^*^Cre^ in the P30 cortex of *Alk4*^fl/fl^ mice indicated that ALK4 is required during embryonic stages for the development of SST^+^ interneuron precursors after these became postmitotic, but prior to the onset of SST expression (Figures 2 and 3). On the other hand, reduced counts of tdTomato^+^ cells in the P30 cortex of *Gad67*^Cre^:*Rosa26*^tdTom^:*Alk4*^fl/fl^ mice indicated a loss of interneurons in the mature cortex of the mutants. In order to determine the timing and possible causes of the loss of SST^+^ interneurons, we assessed earlier stages in the development of these cells in *Gad67*^Cre^:*Alk4*^fl/fl^ mice. At P5, the loss of SST^+^ cells in the cortex of these mice was comparable to that observed at P30, both in its extent and overall layer distribution (Figures 6A and B). At this stage, there was a comparable reduction in tdTomato^+^ cells in layers II-IV and V, indicating cell loss, although the difference in the combined counts over the total cortex did not reach statistic significance due to large value dispersion in the *Gad67*^Cre^:*Alk4*^fl/+^ mice used as controls (Figure 6C). We note that there was no loss of tdTomato^+^ cells in layer VI, despite 50% reduction in SST^+^ cells in this layer (Figures 6B and C), suggesting that P5 may be a transition stage, with loss of SST expression without equivalent loss of cells. As SST protein is not readily detectable in the mouse cerebral cortex before P2 (Forloni et al., 1990), we assessed expression of SATB1, a transcription factor that has been shown to be essential for the development of the SST phenotype in MGE-derived interneurons (Batista-Brito et al., 2008; Close et al., 2012; Denaxa et al., 2012), at earlier stages. To distinguish MGE-derived SATB1^+^ interneurons from other SATB1^+^ cells in the cortex, we assessed the abundance of SATB1/tdTomato double-positive cells in *Gad67*^Cre^:*Rosa26*^tdTom^:*Alk4*^fl/fl^ newborn (P0) mice in comparison to heterozygote controls (*Gad67*^Cre^:*Rosa26*^tdTom^:*Alk4*^fl/+^). We observed a significant decrease in the number of these cells in the mutants (Figures 6D and E), the magnitude of which was generally consistent with the loss of SST^+^ interneurons observed at later stages. However, there was no loss of tdTomato^+^ cells at P0 when comparing homozygote *Alk4*^fl/fl^ mutants to heterozygote *Alk4*^fl/+^ controls (Figure 6F). The small increases observed in the marginal zone (MZ) and layer VI, the two main pathways of tangential migration of cortical GABAergic cells, may have been due to a transient delay in the radial invasion of these cells in the mutants. These data suggest that cell loss begins after birth, in early postnatal stages of *Alk4* mutants. At embryonic stages, SATB1 expression has been detected in postmitotic interneuron precursors from E13.5 onwards (Batista-Brito et al., 2008; Mayer et al., 2018). We examined the number of SATB1^+^ cells in the pallium and subpallium of E14.5 embryos, but found no differences between *Alk4* mutants and controls (Figures 6G to I). In agreement with this, *Satb1* mRNA and protein levels were unchanged in the subpallium of E12.5 and E14.5 *Gad67*^Cre^:*Alk4*^fl/fl^ mutant embryos compared to controls (Figures S7A to C). Finally, we also assessed cell proliferation in the MGE of E12.5 *Gad67*^Cre^:*Alk4*^fl/fl^ and *Nkx2.1*^Cre^:*Alk4*^fl/fl^ embryos, but did not detect any differences when compared to their respective controls (Figures S7D to G), indicating that loss of ALK4 has no impact on cell proliferation in the MGE. In summary, although ALK4 signaling is required during embryonic stages for the correct development of MGE-derived SATB1^+^ precursors of interneurons, cells loss does not begin until the first postnatal week of cortical development in the *Alk4* mutants.

**Figure 6.**
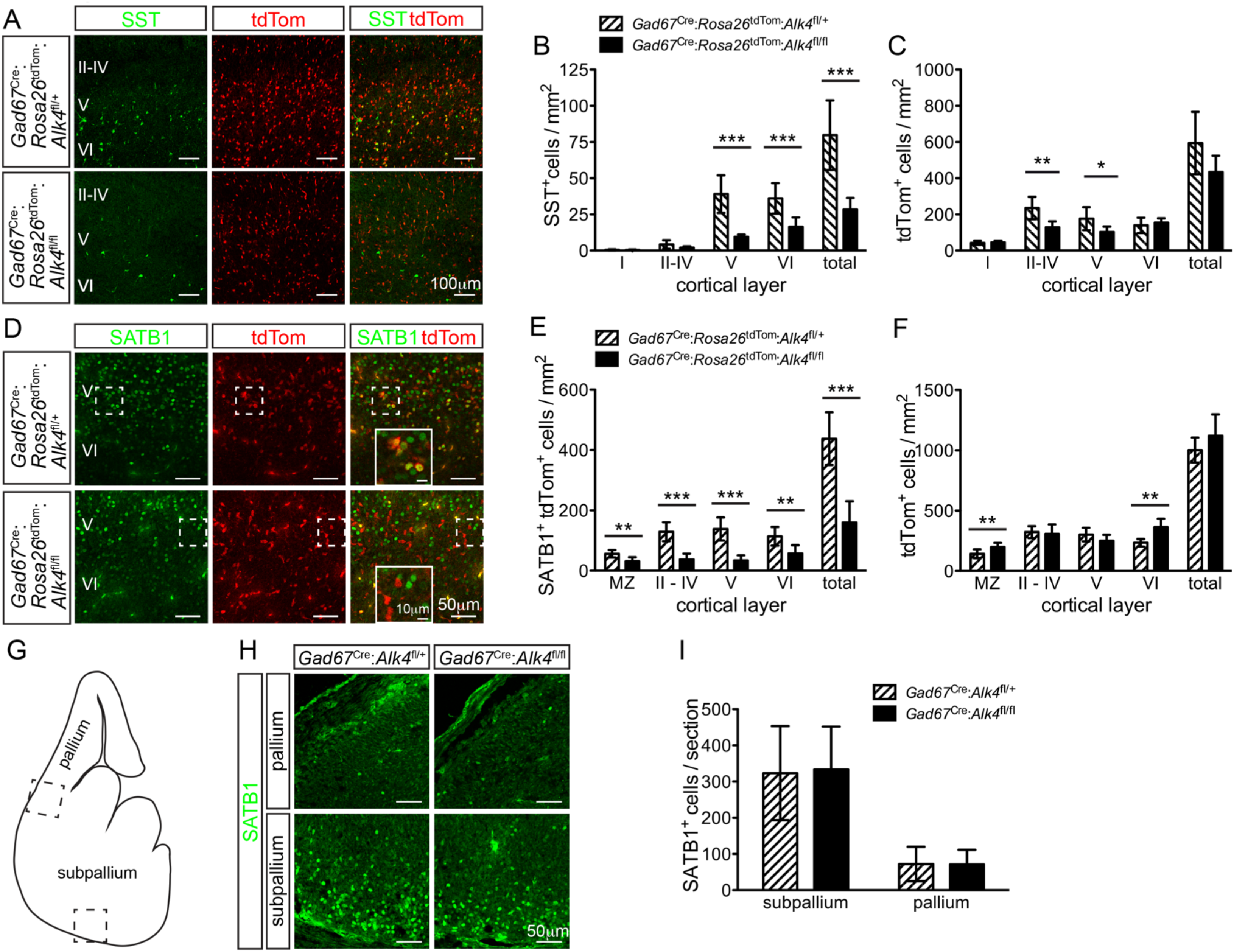
Perinatal loss of SATB1^+^ SST interneuron precursors in *Gad67*^Cre^:*Alk4*^fl/fl^ mutant mice. (A) Representative confocal images of fluorescence immunohistochemical detection of SST (green) and tdTomato fluorescence (red) in the somatosensory cortex of P5 control (*Gad67*^Cre^:*Alk4*^fl/+^, top row) and mutant (*Gad67*^Cre^:*Alk4*^fl/fl^, bottom row) mice. Roman numerals indicate cortical layers as detected by DAPI nuclear counterstaining (not shown here). Scale bar: 100 µm. (B, C) Quantification of SST^+^ (B) and tdTomato^+^ (C) cells in control (*Gad67*^Cre^:*Alk4*^fl/+^, striped bars) and mutant (*Gad67*^Cre^:*Alk4*^fl/fl^, solid bars) mice. Cells were counted in confocal images spanning layers I to VI of the P5 somatosensory cortex. Cell counts were normalized to the area of the section that was imaged and averaged per mouse. Data is presented as mean ± SD. N=6 (control) or 5 (mutant) mice. *, P ≤ 0.05; **, P ≤ 0.01; ***, P ≤ 0.001 (Student’s t-test). (D) Representative confocal images of fluorescence immunohistochemical detection of SATB1 (green) and tdTomato (red) in the cortex of newborn (P0) control (*Gad67*^Cre^:*Rosa26*^tdTom^:*Alk4*^fl/+^, top row) and mutant (*Gad67*^Cre^:*Rosa26*^tdTom^:*Alk4*^fl/fl^, bottom row) mice. Roman numerals indicate cortical layers as detected by DAPI nuclear counterstaining, and layer VI was further verified by TBR1 immunolabeling (not shown here). Insets show magnified views of the areas outlined by the dashed boxes. Scale bars: 50 µm (main images), 10 µm (insets). (E, F) Quantification of SATB1^+^/tdTomato^+^ double-positive (B) and tdTomato^+^ single-positive (C) cells in control (*Gad67*^Cre^:*Rosa26*^tdTom^:*Alk4*^fl/+^, striped bars) and mutant (*Gad67*^Cre^:*Rosa26*^tdTom^:*Alk4*^fl/fl^, solid bars) newborn mice. Cells were counted in confocal images from the marginal zone (MZ) to layer VI of the P0 cortex. Cell counts were normalized to the area of the section that was imaged and averaged per mouse. Data is presented as mean ± SD. N=6 mice per genotype. *, P ≤ 0.05; **, P ≤ 0.01; ***, P ≤ 0.001 (Student’s t-test). (G) Diagram of a coronal section through the E14.5 mouse brain left hemisphere with boxes indicating the approximate location of images shown in (H). (H) Representative confocal images of fluorescence immunohistochemical detection of SATB1 (green) in the pallium and subpallium of control (*Gad67*^Cre^:*Alk4*^fl/+^, left) and mutant (*Gad67*^Cre^:*Alk4*^fl/fl^, right) mice as indicated in (G). Scale bars: 50 µm. (I) Quantification of SATB1^+^ cells in control (*Gad67*^Cre^:*Alk4*^fl/+^, striped bars) and mutant (*Gad67*^Cre^:*Alk4*^fl/fl^, solid bars) E14.5 mouse embryos. Cells were counted in confocal images of the pallium and subpallium in one hemisphere and averaged per mouse. Data is presented as mean ± SD. N=6 mice per genotype. There were no statistically significant differences (P>0.05, Student’s t-test).

### Activin A regulates SATB1 function through ALK4 in MGE cells

Our analysis of SATB1^+^ cells and *Satb1* mRNA and protein levels indicated that the initial expression of this transcription factor is correctly established in the subpallium of *Gad67*^Cre^:*Alk4*^fl/fl^ mutant embryos. We therefore considered the possibility that ALK4 signaling may regulate SATB1 function or its subcellular localization in MGE-derived SST interneuron precursors. In order to assess these possibilities, we established primary cultures of dissociated MGE cells and investigated whether stimulation with activin A, one of the major ligands of ALK4, affected the nuclear localization of SATB1. Activin A induced a significant increase in SATB1 in the nuclei of E12.5 (Figures 7A and B) and E14.5 (Figures S8A and B) MGE cells, with a peak at 30min. Similar results were obtained using biochemical methods in Jurkat cells (Figure S8C), a T cell-derived cell line that is often used in functional studies of SATB1. In MGE cells, the effects of activin A on the nuclear localization of SATB1 were suppressed by the ALK4 inhibitor SB43152 (Figures 7C and D), and activin A had no effect on MGE cells derived from *Gad67*^Cre^:*Alk4*^fl/fl^ mutant embryos (Figures 7E and F). Together, these results suggest that ALK4 signaling may regulate SATB1 function in MGE cells, at least in part, by controlling its nuclear localization.

**Figure 7.**
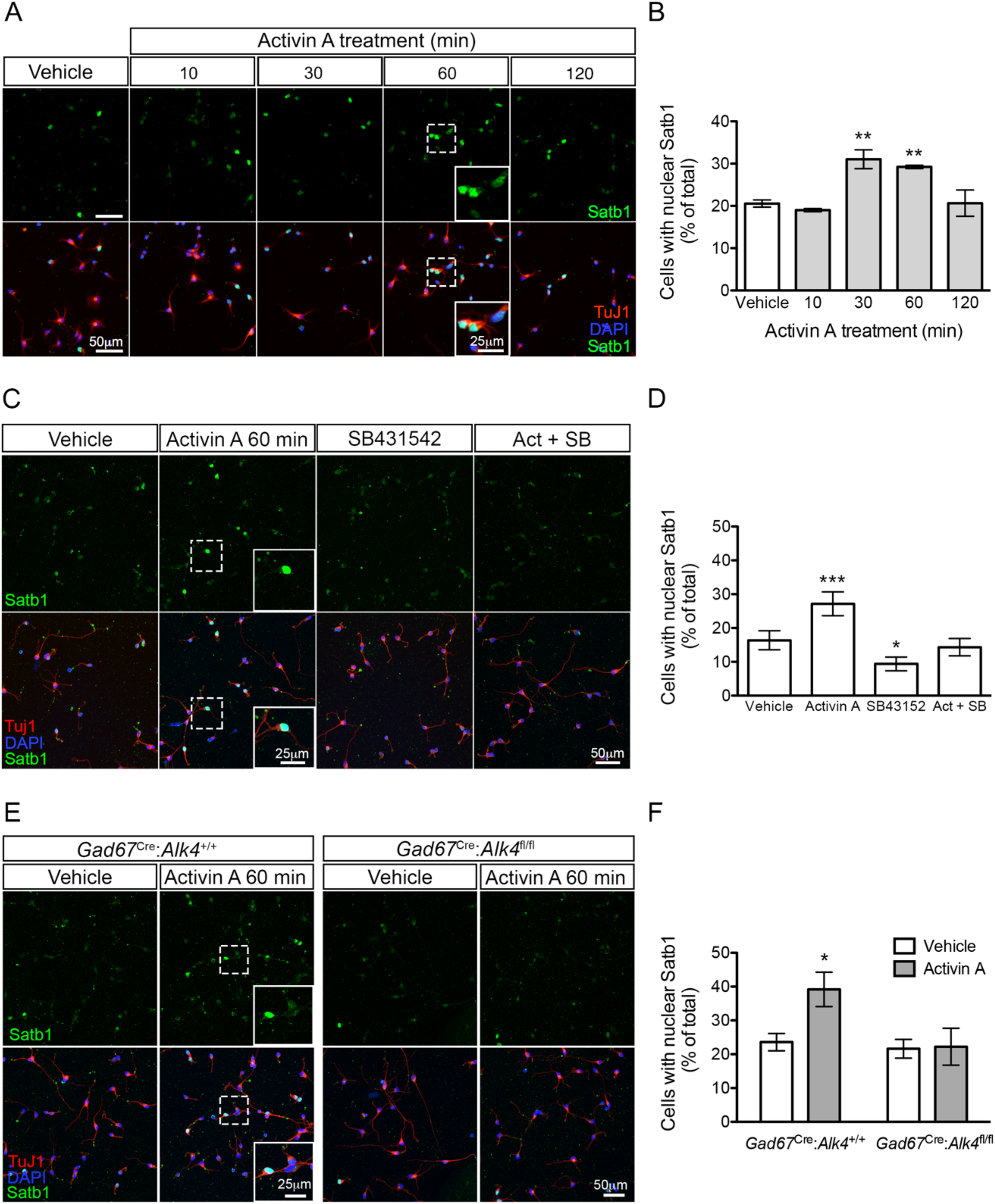
Activin A induces SATB1 nuclear localization in MGE cells. (A) Representative confocal images of fluorescence immunohistochemical detection of SATB1 (green) in cultured MGE cells from wild type E12.5 embryos, counterstained for the neuronal marker Tuj1 (red) and DAPI (blue), after treatment with activin A for the indicated periods of time. Insets show magnified views of areas outlined by dashed boxes. Scale bars: 50µm (main images), 25µm (insets). (B) Quantification of MGE cells displaying nuclear SATB1^+^ after activin A treatment. Results are expressed as percentage of the total cells (mean ± SEM). N=6; **, P < 0.01 compared to vehicle (One-Way ANOVA, P=0.0083, followed by Bonferroni). (C, D) Similar analysis as in (A) and (B) in response to 60 min treatment with activing A or the ALK4 inhibitor compound SB431542 as indicated. N=4; *, P < 0.05; ***, P < 0.001 compared to vehicle (One-Way ANOVA, P<0.001, followed by Bonferroni multiple comparisons test). (E, F) Similar analysis as in (A) and (B) in response to 60 min treatment with activin A on cultured MGE cells derived from control (*Gad67*^Cre^:*Alk4*^+/+^, left) or mutant (*Gad67*^Cre^:*Alk4*^fl/fl^, right) E12.5 embryos as indicated. N=3; *, P<0.05 compared to vehicle (Two-Way ANOVA, P=0.0305, followed by Bonferroni multiple comparisons test).

Activation of type I receptors of the TGFβ superfamily, including ALK4, induces the phosphorylation and nuclear translocation of Smad proteins (Schmierer and Hill, 2007; Budi et al., 2017; Massagué, 2012). We tested the hypothesis that Smad proteins activated by ALK4 may promote the nuclear translocation of SATB1 by direct interaction. To this end, we used the proximity ligation assay (PLA) in cell cultures derived from E12.5 MGE and assessed the interaction between SATB1 and Smad2 under control conditions and in response to activin A. We could detect basal levels of SATB1/Smad2 interaction in control cultures which were significantly increased following 30 min treatment with activin A (Figures 8A and B). Interestingly, a great proportion of the PLA signal in the cultures treated with activin A was found in the cell nucleus (Figure 8A, inset). The SB43152 inhibitor blocked the effect of activin A on SATB1/Smad2 interaction (Figures 8C and D), and activin A had no effect on MGE cells derived from *Gad67*^Cre^:*Alk4*^fl/fl^ mutant embryos (Figures 8E and F). Similar results were obtained using biochemical methods in Jurkat cells (Figure S8D).

**Figure 8.**
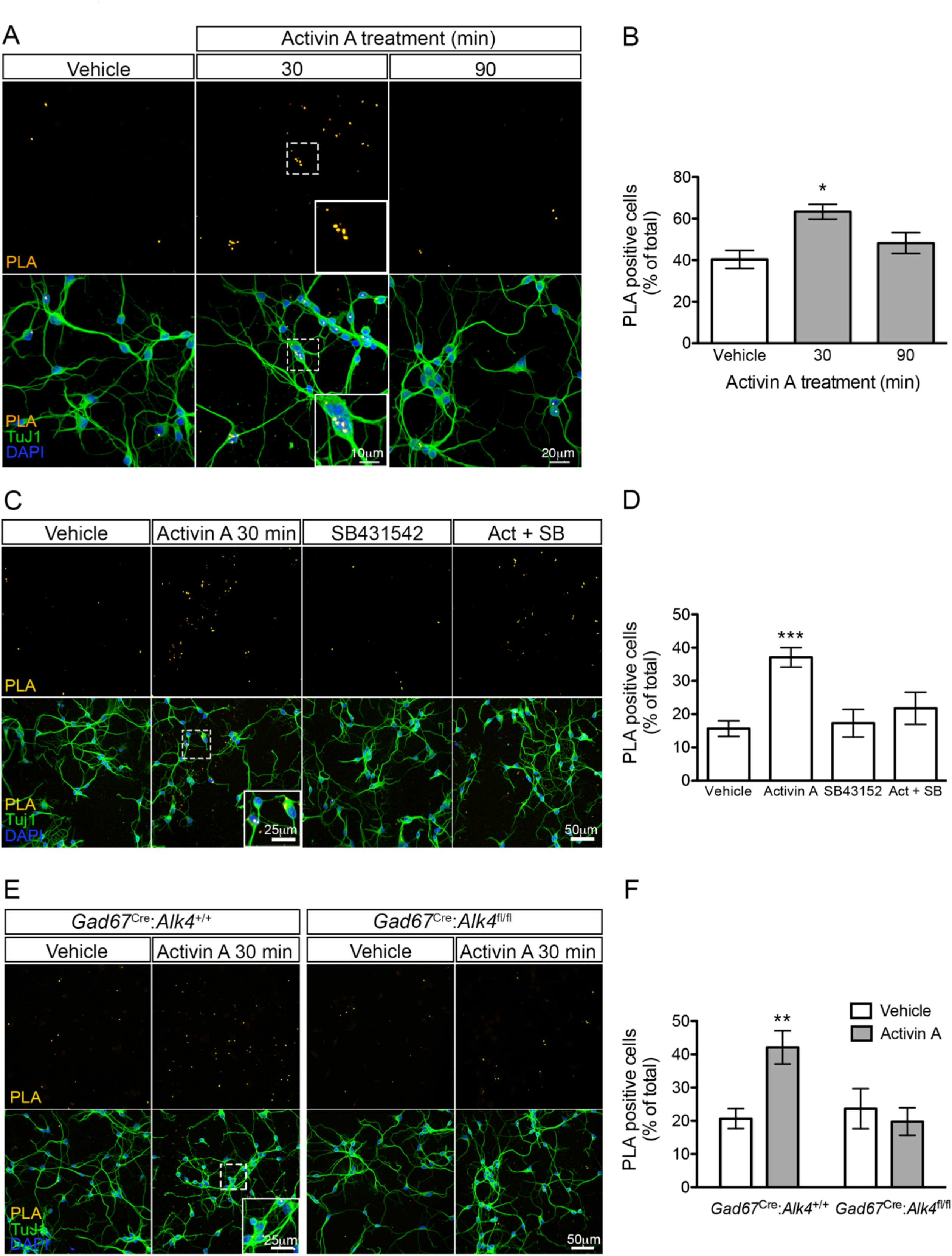
Activin A induces interaction of SATB1 with Smad2 in MGE cells. (A) Representative confocal images of fluorescence immunohistochemical detection of SATB1/Smad2 interaction (PLA, yellow) in cultured MGE cells from wild type E12.5 embryos, counterstained for the neuronal marker Tuj1 (green) and DAPI (blue), after treatment with activin A for the indicated periods of time. Insets show magnified views of areas outlined by dashed boxes. Scale bars: 20µm (main images), 10µm (insets). (B) Quantification of SATB1/Smad2 interaction by PLA in MGE cells after activin A treatment. Results are expressed as percentage of PLA+ cells relative to the total number of cells (mean ± SEM). N=3; *, P<0.05 compared to vehicle (One-Way ANOVA, P=0.01376; followed by Bonferroni multiple comparisons test). (C, D) Similar analysis as in (A) and (B) in response to 30 min treatment with activin A or the ALK4 inhibitor compound SB431542 as indicated. N=4; ***, P<0.001 compared to vehicle (One-Way ANOVA, P<0.001, followed by Bonferroni multiple comparisons test). (E, F) Similar analysis as in (A) and (B) in response to 30 min treatment with activin A on cultured MGE cells derived from control (*Gad67*^Cre^:*Alk4*^+/+^, left) or mutant (*Gad67*^Cre^:*Alk4*^fl/fl^, right) E12.5 embryos as indicated. N=4; *, P<0.01 compared to vehicle (Two-Way ANOVA, P=0.0055, followed by Bonferroni multiple comparisons test).

Finally, we investigated whether the effects of activin A on SATB1 may have a functional significance for SATB1’s role as transcriptional regulator in MGE cells. SATB1 has emerged as a key factor integrating higher-order chromatin architecture with gene regulation. It has also been shown to regulate genes by directly binding to upstream regulatory elements, thereby recruiting chromatin modifiers, corepressors and coactivators directly to gene promoters (Cai et al., 2003; Alvarez et al., 2000; Han et al., 2008). Several reports have shown that SATB1 shows preference for interaction with DNA sequences containing ATC/G triplets (de Belle et al., 1998; Dickinson et al., 1997; Kumar et al., 2005). An enhancer region located 600bp upstream of the transcription start site of the *Sst* gene has been implicated in its regulation by homeobox transcription factors Pdx1 and Pax6 (Andersen et al., 1999). Several ATC/G triplets are located in this enhancer which could potentially serve as binding sites for SATB1. In order to test this notion, we performed chromatin immunoprecipitation (ChIP) assays using antibodies specific for SATB1 as well as control antibodies against CREB, which interacts with a different site in the *Sst* gene promoter (Montminy and Bilezikjian, 1987), and control mouse IgG. We performed PCR amplifications targeting a proximal (R1) and a distal (R2) region, each of about 100bp, within the *Sst* gene enhancer. Both regions could be recovered in SATB1 immunoprecipitates derived from cultures of E12.5 MGE cells (Figures 9A and B). No PCR products could be recovered using antibodies against CREB or control mouse IgG (data not shown). Interestingly, treatment of MGE cells with activin A significantly reduced SATB1 interaction with R1 but increased interaction with R2 (Figures 9A and B), suggesting that activin A signaling through ALK4 can induce relocalization of SATB1 within upstream enhancer sequences in the *Sst* gene of MGE cells.

**Figure 9.**
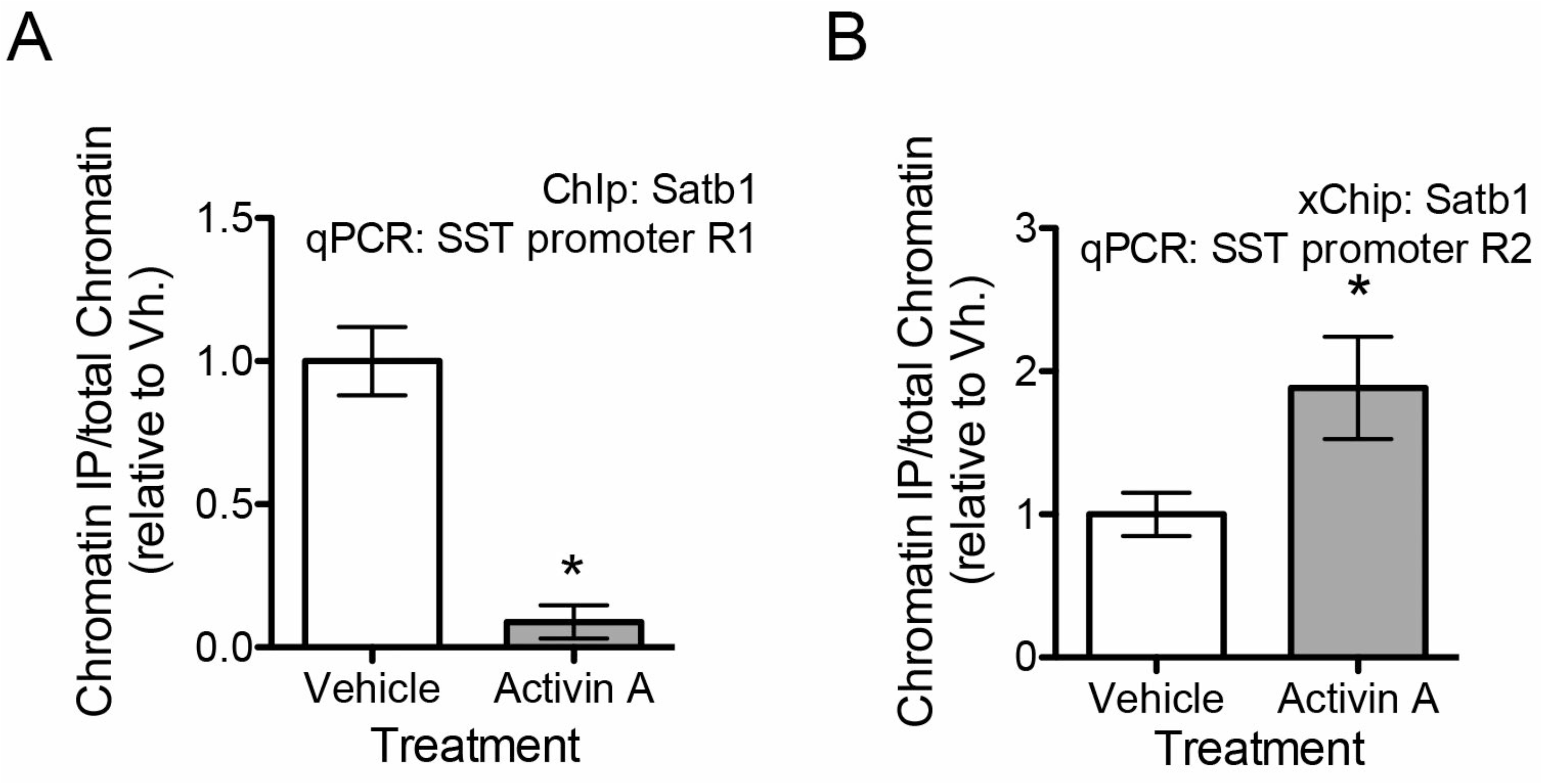
Activin A induces relocation of SATB1 within enhancer regions of the *Sst* gene promoter in MGE cells. (A) Quantification of SATB1 interaction with promoter-proximal R1 enhancer sequence of the *Sst* gene assessed by chromatin immunoprecipitation in MGE cells. Results are presented as mean ± SEM. N= 4*; P< 0.05 compared to vehicle (Student’s t-test, P=0.035). (B) Similar analysis to (A) on promoter-distal R2 enhancer sequence of the *Sst* gene. N= 4*; P<0.05 compared to vehicle (Student’s t-test, p= 0.0431).

## Discussion

Cortical GABAergic interneurons are generated during mouse embryonic development from proliferating progenitors in transient neurogenic zones of the developing basal forebrain. In the MGE, combinations of transcription factors have been suggested to pre-pattern different areas of the VZ and SVZ destined to generate different classes of interneurons (Flames et al., 2007). For example, Nkx2.1 expression marks proliferating progenitors common to both PV^+^ and SST^+^ interneurons (Xu et al., 2008). As they move apically to the mantle zone, these cells leave the cell cycle and begin expression of GABAergic markers, such as GAD67. Topographical transcriptome analysis of this region revealed a progression in the expression of subsets of genes implicated in interneuron differentiation, neuronal migration and neuronal projection, as the cells move ventro-laterally (Zechel et al., 2014). From the MGE, GABAergic cells migrate tangentially to colonize the developing cortex and hippocampus (Corbin et al., 2001; Marín and Rubenstein, 2001; Wichterle et al., 2001). Throughout these processes, extracellular factors have so far been mainly implicated in the guidance of migrating GABAergic cells from the subpallium to the pallium and in the developing neocortex (e.g. (Flames et al., 2004; Pozas and Ibáñez, 2005; Lopez-Bendito et al., 2008)). On the other hand, whether extracellular signals interact with intrinsic transcriptional programs to promote the specification of interneuron subtypes in the subpallium or beyond, remains less well understood. The studies presented here implicate activing signaling through the type I TGFβ superfamily receptor ALK4 in the development of SST^+^ interneurons through regulation of the intracellular localization and activity of SATB1, a key transcriptional regulator of the differentiation and maturation of these cells. ALK4 was expressed in Lhx6^+^ cells of the MGE during the peak of SST^+^ interneuron generation along with several of its ligands. The effects of the ALK4 inhibitor SB43152 on SATB1 nuclear localization on cultured neurons derived from the MGE also suggests endogenous production of ligands by these cells. ALK4 could therefore be part of a quorum-sensing mechanism ensuring that appropriate numbers of SST interneuron precursors are generated during embryonic development.

The loss of tdTomato^+^ cells in the cortex of *Gad67*^Cre^:*Rosa26*^tdTom^:*Alk4*^fl/fl^ mice at P30 and P5 indicates a loss of interneurons, rather than mere downregulation of marker gene expression. At P30, this loss was quantitatively comparable to the combined loss observed in non-overlapping subpopulations of PV^+^, SST^+^ and SST ^-^/RELN^+^ interneurons, indicating that these are all the major cell types affected by the loss of ALK4 in GABAergic cells of the developing basal forebrain. At P5, however, the loss of tdTomato^+^ cells was lower than that expected from the SST^+^ counts, particularly in layer VI, suggesting that this stage may be close to the onset of cell elimination in the mutants. Although the *Sst* gene initiates transcription during the later stages of embryonic development (Batista-Brito et al., 2008), SST protein is not detected until the first postnatal days in the mouse brain (Forloni et al., 1990). To visualize SST interneuron precursors, we used SATB1 as a prospective marker of these cells. We note that, although SATB1 is also expressed by PV^+^ cells, the latter population was not as significantly affected by the loss of ALK4 as the SST^+^ population when analyzed in the adult brain (compare Figures 2D and 2E). Therefore, we believe that the vast majority of changes detected perinatally in SATB1^+^ /tdTom^+^ cells should represent changes of the SST interneurons. While the loss of SATB1^+^ cells at birth was quantitatively consistent with the extent of SST^+^ neuron loss observed later in the cortex, we did not detect a comparable loss of tdTomato^+^ cells at this stage, indicating that phenotypic defects precede cell elimination in the *Alk4* mutant cortex. At E14.5, no loss of SATB1^+^ cells could be detected in the pallium or subpallium of the mutants. In addition, ALK4 was dispensable for the proliferation of VZ and SVZ progenitors of these cells, and it was also dispensable for the maintenance of the SST phenotype, as *Sst^IRES-^*^Cre^:*Alk4*^fl/fl^ mice showed no loss of SST^+^ interneurons. Based on these observations, we conclude that ALK4 is required for the correct development of SST interneuron precursors during a relatively narrow time window of embryonic development, after these cells become postmitotic and GABAergic, but before they activate the *Sst* locus itself a few days later. Lack of ALK4 signaling during this critical period resulted in impaired differentiation at later stages, as shown by an inability to maintain normal SATB1 expression at birth, and eventually cell elimination during the first postnatal week. Many factors are known to act transiently during early stages to influence phenotypes that manifest later in development. We note that the timing of cortical SATB1^+^/SST^+^ cell elimination in the *Alk4* mutants approximately coincided with the period of intrinsic elimination of cortical GABAergic cells identified by recent studies, which was reported to occur between P3 and P13, with a marked peak at P7 (Southwell et al., 2012). It is possible that this intrinsic cell death may serve to eliminate GABAergic cells that failed to differentiate properly, as in the case of our *Alk4* mutants.

Unlike several other interneuron classes, SST^+^ cells in cortical layer IV, also referred to as X94 cells, target PV^+^ fast-spiking interneurons in the same layer (Xu et al., 2013). The morphology of these cells also differs from other SST^+^ interneurons, as they are often described as bitufted or multipolar cells with local axonal projections (Ma et al., 2006; Yavorska and Wehr, 2016). *Alk4* deletion driven by either *Gad67*^Cre^ or *Nkx2.1*^Cre^ resulted in the loss of about two-thirds of SST^+^ cells in layer IV, and approximately half of the cells expressing *Hpse* mRNA, now considered a specific marker of X94 cells (Naka et al., 2018). The significant loss of X94 cells in the *Alk4* mutants correlated with the reduction in both the amplitude and frequency of IPSCs in PV^+^ interneurons in the same layer. Although it has been shown that fast spiking PV ^+^ interneurons can inhibit other PV^+^ cells in this layer, the moderate decrease in PV^+^ neurons that we detected in *Alk4* mutants (statistically significant only with respect to WT layer IV, but not in comparison to all other controls), would seem insufficient to account for the substantial effect observed on inhibition, which was greater than 75%. Based on these results, we conclude that X94 cells are a subpopulation of SST/HPSE double positive interneurons that depend on ALK4 for their correct development. Because of the early requirement of ALK4 for the development of these cells, our results provide functional evidence for the pre-specification of the X94 phenotype during embryonic stages. A Martinotti subtype of layer V SST^+^ interneurons that project to layer I was completely ablated in the *Alk4* mutants, indicating their absolute dependence on ALK4 signaling during early embryonic stages. On the other hand, no loss of *Chodl* mRNA^+^ cells could be detected, indicating that not all subtypes of cortical SST^+^ neurons depend on ALK4 for their correct development. Intriguingly, these cells are in fact long-projecting GABAergic neurons, not properly interneurons, a fact that may be related to their ALK4 independence. Together, these results suggest that dependence on ALK4 is not stochastic among developing precursors of SST interneurons, but rather that some subtypes have a strict dependence on ALK4 for their correct specification, while others are totally independent.

SATB1 has been shown to be essential for the specification of the SST interneuron phenotype (Batista-Brito et al., 2008; Close et al., 2012; Denaxa:2012bk; Narboux-Nême et al., 2012). SATB1 can affect gene expression at multiple levels. It can tether different genomic loci to help coordinate their expression, recruit chromatin-remodeling enzymes to regulate chromatin structure, and interact directly with gene promoter and enhancer regions to engage co-activators or co-repressors (Alvarez et al., 2000; Cai et al., 2003; Kumar et al., 2006). Our finding of an interaction between Smad proteins and SATB1 brings a new perspective on the functions of this genome organizer and transcriptional regulator, linking its activity to information carried by extracellular signals. SATB1 bears an atypical nuclear localization signal (Nakayama et al., 2005) and the regulation of SATB1 subcellular localization is not well understood. A proteomics study found altered levels of SATB1 in the nuclear fraction of human CD4^+^ T cells upon stimulation with IL-4 (Moulder et al., 2010), suggesting that the nuclear localization of SATB1 is indeed susceptible to regulation by extracellular stimuli, although the mechanisms involved were not elucidated. We found that activin signaling could increase the nuclear localization of SATB1 in MGE-derived neurons, and PLA studies indicated that SATB1 could be found in complex with Smad2 in the nucleus of these cells, suggesting that activated Smad proteins may bring along SATB1 as they shuttle to the cell nucleus. Using chromatin immunoprecipitation, we found that activin signaling altered the interaction of SATB1 within the enhancer region of the *Sst* gene in MGE cells, decreasing its binding to a distal region while increasing it to a more proximal region. This result suggests that, through SATB1, activin signaling contributes to reorganize the *Sst* locus in preparation for initiation of gene transcription. We note, however, that *Sst* mRNA levels were not increased in cultured MGE cells stimulated with activin A, suggesting that activin signaling is not itself sufficient to regulate *Sst* gene transcription.

In summary, the results of the present study reveal a previously unknown level of regulation in the specification of cortical interneuron subtypes mediated by an extracellular signal during early embryonic development. It is likely that many other transcriptional programs affecting cortical interneuron diversification also interact with extracellular signals in the environment of the ganglionic eminences and along the migratory routes of interneuron precursors to coordinate GABAergic interneuron development in the mammalian brain.

## Summary of online supplementary materials

Online supplementary materials include Supplementary Methods for Jurkat cell culture, Western blotting, immunoprecipitation and nuclear fractionation; Supplementary Figure S1 to S5 and Supplementary table S1.

## Methods

### Mice

Mice were housed in a 12 hour light-dark cycle, and fed a standard chow diet. The *Alk4*^fx^ knock-in allele was generated by introducing LoxP sites in the IVth and VIth introns of the *Acvr1b* gene, encoding ALK4 (Figure S1). Targeting vectors were generated using BAC clones from the C57BL/6J RPCIB-731 BAC library and transfected into TaconicArtemis C57BL/6N Tac ES cell line. Gene-targeted mice were generated at TaconicArtemis by standard methods. Both female and male mice were used in the study. The day of vaginal plug was considered embryonic day 0.5 (E0.5). All animal procedures are in accordance with Karolinska Institute’s ethical guidelines and were approved by Stockholms Norra Djurförsöksetiska Nämnd.

### Antibodies

The primary antibodies used for immunohistochemistry were as follows: mouse anti-PV (1:1000, 235, Swant), rabbit anti-SST (1:2000, T-4103, Peninsula), mouse anti-RELN (1:1000, MAB5364, Millipore), rabbit anti-VIP (1:500, 20077, Immunostar), goat anti-SATB1 (1:100, sc5889, Santa Cruz), anti-RFP (1:500, 600-401-379, Rockland), and rat anti-BrdU (1:500, OBT0030G, Accurate Chemicals). The secondary antibodies used for immunohistochemistry, all raised in donkey and used at 1:2000 dilution, were as follows: anti-mouse Alexa Fluor 488 (A21202, Invitrogen), anti-mouse Alexa Fluor 555 (A31571, Invitrogen), anti-rabbit Alexa Fluor 555 (A31572, Invitrogen), anti-goat Alexa Fluor 488 (A11055, Invitrogen), anti-rat Alexa Fluor 647 (712-606-153, Jackson Immunoresearch), and anti-rat Alexa Fluor 488 (A21208, Invitrogen).

### Tissue preparation, fluorescence immunohistochemistry and in situ hybridization

Male and female postnatal day 30 (P30) mice were deeply anaesthetized, transcardially perfused with PBS and 4% PFA in PBS, and brains were dissected and postfixed overnight at 4°C. 40 µm free-floating coronal sections were cut at the vibratome (VTS 1200, Leica, Germany) and subjected to fluorescence immunohistochemistry: blocking, permeabilization and antibody incubation were carried out in 5 % normal donkey serum (NDS; Jackson Immunoresearch), 0.2% Tx-100 in PBS. Primary antibodies were incubated overnight at 4°C, secondary antibodies for 2 h at room temperature (RT). Nuclei were visualized with DAPI. All washes were performed with PBS, pH 7.4. Sections were mounted on object slides, air-dried and coverslipped using DAKO fluorescence mounting medium (Dako North America). P5 mice were anaesthetized, transcardially perfused with PBS and their brains were dissected out and fixed overnight in 4 % PFA. P0 mice and embryos were quickly decapitated the heads rinsed in PBS and immersed in 4% PFA (6 h – overnight). Brains were cryoprotected in 30 % sucrose and embedded in OCT cryomount (Histolab products AB, Sweden) for cryostat sectioning. Coronal sections (16 µm, series of 10 for P5 and P0 mice; 12 µm, series of 8 for embryos) were thaw-mounted onto Superfrost^+^ slides (Menzel Gläser, Germany), air dried, stored at −80°C and subjected to immunohistochemistry as described below. To detect BrdU sections were incubated in 1M HCl for 45 min at 37°C followed by 15 min at RT in 10mM Tris-HCl pH 8 and washes in PBS prior to the general protocol outlined above. Detection of SATB1 required heat mediated antigen retrieval in 10mM citric acid buffer pH 6 prior to the general immunohistochemical procedure. For RNAscope embryos were rapidly decapitated and the heads were fresh-frozen in OCT and stored at −80° C. Adult mice were anaesthetized, decapitated and their brains were dissected out for freezing in OCT and storage at −80° C. Coronal cryostat sections (10µm thick) were thaw-mounted onto Superfrost^+^ slides and stored at −80° C until further processing. In situ hybridization was carried out using RNAscope technology (Advanced Cell Diagnostics Biotechne), following the manufacturer’s protocol. The following probes were utilized: Alk4 (Mm-Acvr1b, Cat No. 429271; Mm-Lhx6-C3, Cat No.422791-C3).

### Image acquisition and analysis

Images were acquired using a LSM700 confocal microscope (Zeiss, Germany). To analyze the expression of interneuron markers at P30, a 15.86µm confocal stack was imaged in the somatosensory cortex from every sixth section between Bregma 1.94 mm to −2.18 mm. At P5 and P0 corresponding regions were imaged. Embryonic sections containing both MGE and LGE were imaged using the tile scan function. To analyze recombination efficiency images from three non-adjacent sections from the somatosensory cortex were acquired. Cells were counted using the Cell Counter plugin in Image J (NIH Image J 1.51), counts are reported as mean ±SD. Statistical data analysis was performed using GraphPad Prism 5.

### Quantitative real-time PCR (qRT-PCR)

E12.5 and E14.5 control and mutant embryos were decapitated and their brains dissected in ice-cold sterile PBS supplemented with 1% glucose. MGE and LGE were frozen on dry-ice in Eppendorf tubes and stored at −80°C. RNA extraction and cDNA synthesis were performed in parallel for all samples. Total RNA was extracted using the RNeasy Mini Kit (Qiagen, Hilden, Germany) according to the manufacturer’s protocol. Subsequent to a DNaseI digest (Invitrogen ThermoFisherScientific, Waltham, USA) cDNA was synthesized from 250µg of total RNA primed with random hexamers using SuperScript II (Invitrogen). qRT-PCR analysis was performed using SYBR green PCR Master Mix (Applied Biosystems, Foster City, USA) according to the manufacturer’s protocol on a StepOnePlus continuous fluorescence detector (Applied Biosystems) under standard cycling conditions. The oligonucleotides used were as follows: Gdf1 Rv: AGGTCAAAGACGACTGTCCA; InhbA Fw: ATCATCACCTTTGCCGAGTC; InhbA Rv: ACAGGTCACTGCCTTCCTTG; InhbB Fw: CTTCGTCTCTAATGAAGGCAACC; InhbB Rv: CTCCACCACATTCCACCTGTC; Satb1 Fw: ACAGTAAGGAATGCTCTGAAGG, Satb1 Rv: CTGTTCACAATGGAGGAGATCA; 18S Fw: CACACGCTGAGCCAGTCAGT; 18S Rv: AGGTTTGTGATGCCCTTAGATGTC (Fw, forward; Rv, reverse). Formation of specific amplicons was verified by melt curve analysis, gene expression was quantified relative to 18S ribosomal RNA.

### Acute slice electrophysiology

Postnatal day 15 to 19 *Nkx2.1*^Cre^:*Rosa26*^tdTom^:*Alk4*^f*l/+*^ and *Nkx2.1*^Cre^:*Rosa26*^tdTom^:*Alk4*^fl/fl^ pups were anesthetized and brains were collected in ice-cold solution of the following composition (in mM): 62.5 NaCl, 100 sucrose, 2.5 KCl, 25 NaHCO_3_, 1.25 NaH_2_PO_4_, 7 MgSO_4_, 1 CaCl_2_, and 10 glucose. Subsequently, brains were sectioned to 300µm slices, which were then let recover for 1 h at room temperature in oxygenated aCSF (in mM): 125 NaCl, 2.5 KCl, 25 NaHCO_3_, 1.25 NaH_2_PO_4_, 2 MgSO_4_, 2 CaCl_2_, and 10 glucose. During whole-cell patch-clamp recordings, slices were perfused with oxygenated aCSF at 24±2°C. Patch electrodes were made from borosilicate glass (resistance 4–8 MΩ; Hilgenberg, GmbH) and filled with a solution containing (in mM): 65 K-gluconate, 65 KCl, 10 HEPES, 10 Na-Phosphocreatine, 0.5 EGTA, 4 MgATP, 0.3 Na2GTP. We targeted layer 4 cells in the somatosensory cortex, either tdTomato-positive (to enrich for PV-positive fast-spiking neurons). Intrinsic properties of cells were recorded in current-clamp and analyzed as previously described (Muñoz-Manchado et al., 2018). In our conditions, the Cl^-^ reverse potential determines an outward current of this ion when GABA receptors are open, shown in the recordings as excitatory currents. In order to measure inhibitory post-synaptic currents (IPSCs), neurons were recorded in voltage-clamp mode held at −70 mV in aCSF containing glutamatergic blockers CNQX/MK-801 (10µM/5µM, Sigma) for at least 10 min. Currents were recorded with an Axopatch 200B amplifier (Molecular Devices), sampled at 10 kHz, processed with Digidata 1322A (Molecular Devices) and analyzed on Clampex. Traces were low-pass filtered (1KHz), only events with amplitude larger than −10pA were included in the analysis, and frequency was calculated as total number of events per time. Data are mean ± SEM. P values are from Student’s two-tailed t-tests.

### Cell culture

For primary culture of MGE cells, brains from E12.5 or E14.5 mouse embryos were collected and placed in ice-cold PBS supplemented with 1% glucose. The MGE was dissected out and digested with EDTA-Trypsin buffer (Gibco, Thermofisher) during 15min. Trypsin was inactivated with 25% Fetal Bovine Serum (FBS, Gibco), and cells were dissociated through a glass pipet in the presence of 25ng DNase (Roche). Cells were collected by centrifugation during 5 min at 800g and resuspended in Neurobasal Medium (Gibco) supplemented with B27 (Gibco), 2mM Glutamine (Gibco) and 20µg/ml Penicilin/stretptomicyn. MGE cells were plated at a density of 50,000 cells per coverslip in a 24-well plate or, alternatively, at 150,000 cells per well in 12 wells-plates; both coated with 100µg/ml Poly-D-Lysine and 2µg/ml of Laminin (Cultrex, RnD). Cells were maintained during two days at 37°C in 95% O_2_/5% CO_2_ atmosphere before the start of the experiments.

Jurkat cells were growth in RPMI-1640 medium containing 2mM glutamine (Gibco) 20ug/ml Penicillin/streptomycin (Gibco) and 10% Fetal Bovine Serum (Gibco), at 37°C in 95% O_2_/5% CO_2_ atmosphere.

### Satb1 nuclear translocation and proximity ligation assays in MGE cells

After 2 days in culture, MGE cell monolayers were treated with activin A at 100ng/ml (RnD) for the indicated periods of time. In some experiments, the ALK4 inhibitor SB41542 (Sigma-Aldrich) was added at 10µM 60min prior to the addition of activin A, or on its own. At the end of the treatment, cells were washed twice with PBS and fixed for 15 min in 4% paraformaldehyde (PFA)/4% sucrose, permeabilized, and blocked in 10% normal donkey serum (NDS) and 0.3% Triton X-100 in PBS. Cells were then incubated overnight at 4°C with anti-SATB1 (Santa Cruz; sc5989; 1:400), and anti-TuJ1 (βIII Tubulin, Sigma-Aldrich, MAB1637;1:2000) in PBS supplemented with 5% NDS and 0.05% Triton. After washing with PBS, the cultures were incubated with the appropriate secondary antibody. Secondary antibodies were Alexa-Fluor-conjugated anti-immunoglobulin from Life Technologies and Invitrogen, used at 1:1000 (donkey anti-rabbit immunoglobulin G [IgG] Alexa Fluor 555, A31572; donkey anti-goat IgG Alexa Fluor 488, A11055; donkey anti-mouse IgG Alexa Fluor 488, A21202; donkey anti-mouse IgG Alexa Fluor 555, A31570; donkey anti-mouse IgG Alexa Fluor 647, A31571; donkey anti-goat IgG Alexa Fluor 555, A21432). For proximity ligation assay (PLA), cells were fixed as above and incubated overnight at 4°C with anti-SATB1 (Santa Cruz; sc5989, 1:200), anti-Smad2 (Cell Signalling; 5339; 1:100), and anti-Tuj1 (Sigma-Aldrich; MAB1637;1:2000) antibodies in PBS supplemented with 3% BSA. The Duolink *In Situ* Proximity Ligation kit (Sigma) was then used as per manufacturer’s instructions with fluorophore-conjugated secondary antibody to recognize Tuj1 (donkey anti-mouse IgG Alexa Fluor 488, A21202, 1:1000) included during the amplification step. Cells were imaged with an Zeiss LSM700 confocal laser microscope. Image analysis was performed using NIH Image J 1.51 and GraphPad Prism 5.

### Western blotting, immunoprecipitation and nuclear fractionation

Protein samples of Jurkat cells were prepared for SDS-PAGE in SDS sample buffer (Life Technologies). Protein samples of MGE tissue were lysed in RIPA buffer containing protease inhibitor (Roche) using a 25G syringe. Total protein concentration was measured using a Dc-Protein Assay (Bio-Rad). 25µg of protein were loaded per sample Protein samples were boiled at 95°C for 10 min before electrophoresis 12% polyacrylamide gels. Proteins were transferred to polyvinylidene fluoride (PVDF) membranes (Amersham). Membranes were blocked with 5% non-fat milk and incubated with primary antibodies. The primary antibodies used for Western blotting and immunoprecipitation were as follows: goat anti-SATB1 (Santa Cruz; 5889; 1:600); rabbit anti-Smad2 (Cell Signaling; 5339; 1:1500); and mouse anti-αTubulin (Sigma-Aldrich; T6199; 1:2000). Horseradish peroxidase (HRP)-conjugated secondary antibodies for Western blotting were from DAKO (Agilent). Immunoblots were developed using Clarity Western ECL (BioRad) or SuperSignal West Femto Maximum Sensitivity Substrate (ThermoFisher). Images were acquired through a LAS 4000-ImageQuant System (GE Healthcare); quantification of band intensities was done with ImageQuant software (GE Healthcare). For immunoprecipitation, cells were lysed with RIPA buffer containing protease inhibitor (Roche). Total protein was collected and incubated with anti-Satb1 antibody (Santa Cruz; 5889; 1:50) overnight at 4°C and then incubated with SepharoseProtein-G beads (GE Healthcare). Samples were then prepared for immunoblotting as described above. For nuclear/cytoplasmic fractionation, nuclear and cytosolic fractios were collected using the NE-PER™ Nuclear and Cytoplasmic Extraction Reagents (ThermoFisher) according to the manufacturer protocol.

### Chromatin immunoprecipitation

MGE cells treated with Activin A (100ng/ml) during 60 minutes, collected by trypsinization, and washed twice in ice cold-PBS. Cells of 3 wells from 12 well-plates were pooled together (around 450,000 cells in total). DNA-protein complexes were crosslinked by incubating the cells with 37% formaldehyde (final concentration 1%) during 8 minutes. DNA was then purified and immunoprecipitated using a HighCell ChIP Kit (Diagenode, Belgium) according to the manufacturer’s protocol. DNA was sheared by sonication using a Sonicator (Diagenode, Belgium): 10 cycles (30 seconds ON, 30 seconds OF) at high power setting. DNA was immunoprecipitated using an anti-SATB1 antibody (Santa Cruz, X-5889, 6.3 µg/sample), or anti-CREB (Abcam 31387; 6.3µg/sample) or mouse-IgG (provided with the kit). Immunoprecipitates were amplified and analyzed by real-time qPCR using SybrGreen reagents (Applied Biosystems). The following qPCR primers were used: SST-promoter-R1: Fw: CCTGTAGGGATCATCTCGTCC; Rv: GGCCAGAGTTCTGACTGCTTT. SST-promoter-R2: Fw ACTCTGGCCTGAACAGTAAACAT; Rv: TCAGCTCTGCCTGATCTCCTA. IL2a-promoter: Fw GGGGGTGGGGATACAAAGTAA. Rv: TCTTGCTCTTGTCCACCACAATA.

### Statistical analysis

Statistics analyses were performed using Prism 5 software (GraphPad, SPSS IBM corporation) and Microsoft Excel (Microsoft). Student’s t test, one-way ANOVA or two-way ANOVA were performed to test statistical significance according the requirements of the experiment (e.g. comparisons across one or two variables, respectively). Bonferroni test was used as a further analysis for experiments that required multiple comparisons. Statistical analysis of cumulative curves was done with the Unpaired Kologorov-Smirnov test, which tests the similarity of continuous probability distributions, without any assumptions on data distribution, and is often used to analyze cumulative curves. Differences were considered statistically significant when P<0.05. All P values are reported in Table S1.

## Acknowledgements

We thank Wei Wang for technical assistance, and Victor Tarabykin and Gord Fishell for antibody and tissue used in pilot studies. Support for this research was provided by grants to C.F.I. from the Swedish Research Council (2016-01538), Knut and Alice Wallenbergs Foundation (KAW 2012.0270), and the National University of Singapore (R-185-000-227-133); and to J. H.-L. from Swedish Research Council (2014-3863), StratNeuro, and the Swedish Brain Foundation. The authors declare no conflict of interests.

## Author contributions

C.G. performed the studies shown in Figures 2, 3, 4, 6, S2 and S3. F.A.K. performed the studies shown in Figures 1, 7, 8, 9, S1B, S4 and S5. C.G. and F.A.K. should be considered as equal first authors. H.M. performed the studies shown in Figure 5. D.F.S. and A.A. provided assistance with mouse breeding, genotyping and histological studies. C.G., F.A.K., H.M., J.H.-L. and C.F.I. performed data analysis. C.G. wrote a first draft and C.F.I. wrote the final paper.

---

**Figure S1.**
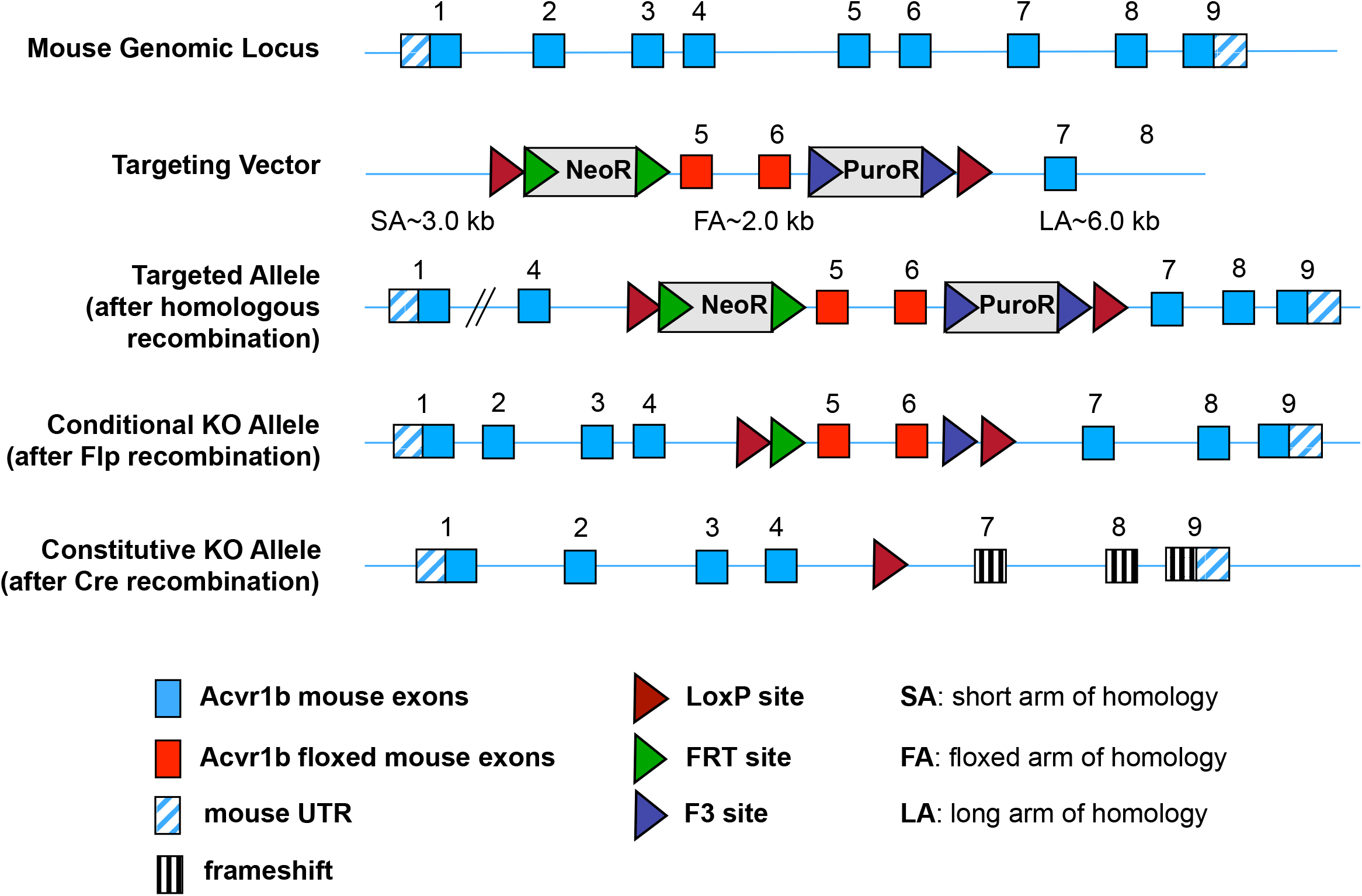
Generation of a conditional allele of the mouse *Acvr1b* gene encoding ALK4. CRE-mediated recombination deletes exons 5 and 6, encoding the ALK4 kinase domain, which is essential for signaling, and introduces an in-frame stop codon after exon 4.

**Figure S2.**
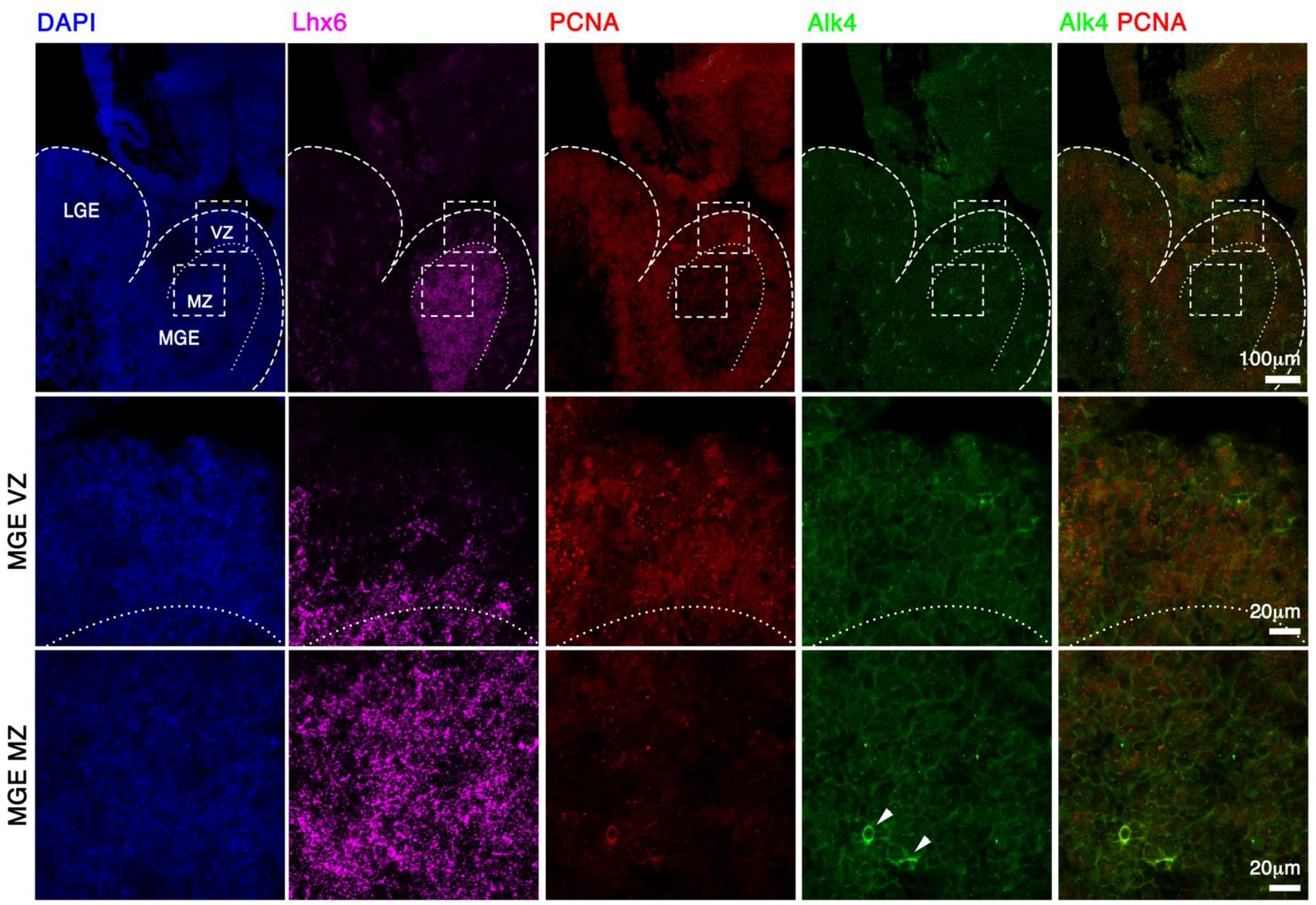
Analysis of *Alk4* mRNA expression in the MGE of *Alk4* mutant mice. RNAscope in situ hybridization analysis of *Alk4* (green) and *Lhx6* (purple) mRNA and PCNA protein (red) expression in mutant (Gad67Cre:Alk4fl/fl) E12.5 mouse embryos. Middle and lower rows show higher magnification images of ventricular (VZ) and mantle **(MZ)** zones, respectively, of areas boxed in upper panel. *Alk4* mRNA expression was abolished in areas positive for Lhx6 mRNA in these mutants (compare to wild type shown in Figure 1). Arrowheads point to blood vessels (showing unspecific signal in the green channel). Scale bars: 100µm (upper row); 20µm (middle and lower rows).

**Figure S3.**
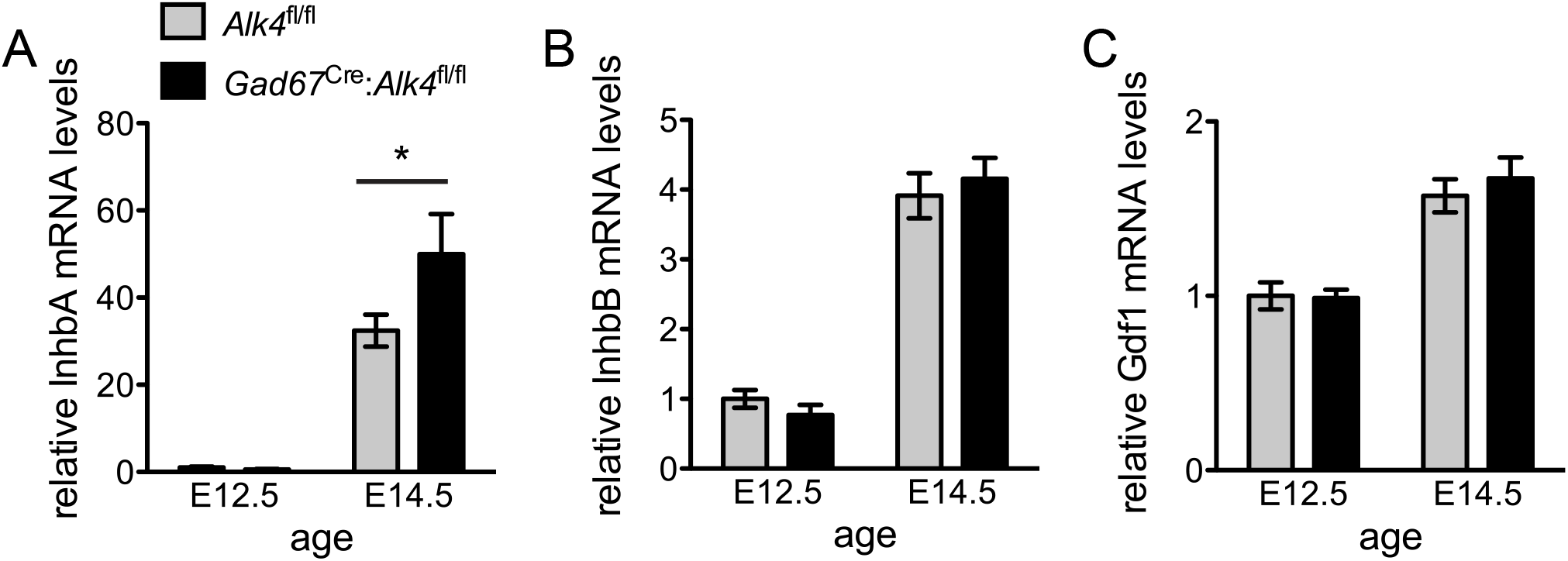
Expression of mRNAs encoding ALK4 ligands in the embryonic basal forebrain. Expression of Inhba (A), Inhbb (B) and Gdf1 (C) mRNAs in basal forebrain of control (Alk4fl/fl, grey bars) and mutant (Gad67Cre:Alk4fl/fl, black bars) E12.5 and E14.5 mouse embryos. Results are presented as average ± SEM. N=8 (E12.5 Alk4fl/fl), 6 (E12.5 Gad67Cre:Alk4fl/fl), 6 (E14.5 Alk4fl/fl), 8 (E14.5 Gad67Cre:Alk4fl/fl). *, P < 0.05 (ANOVA, Bonferroni post-hoc test).

**Figure S4.**
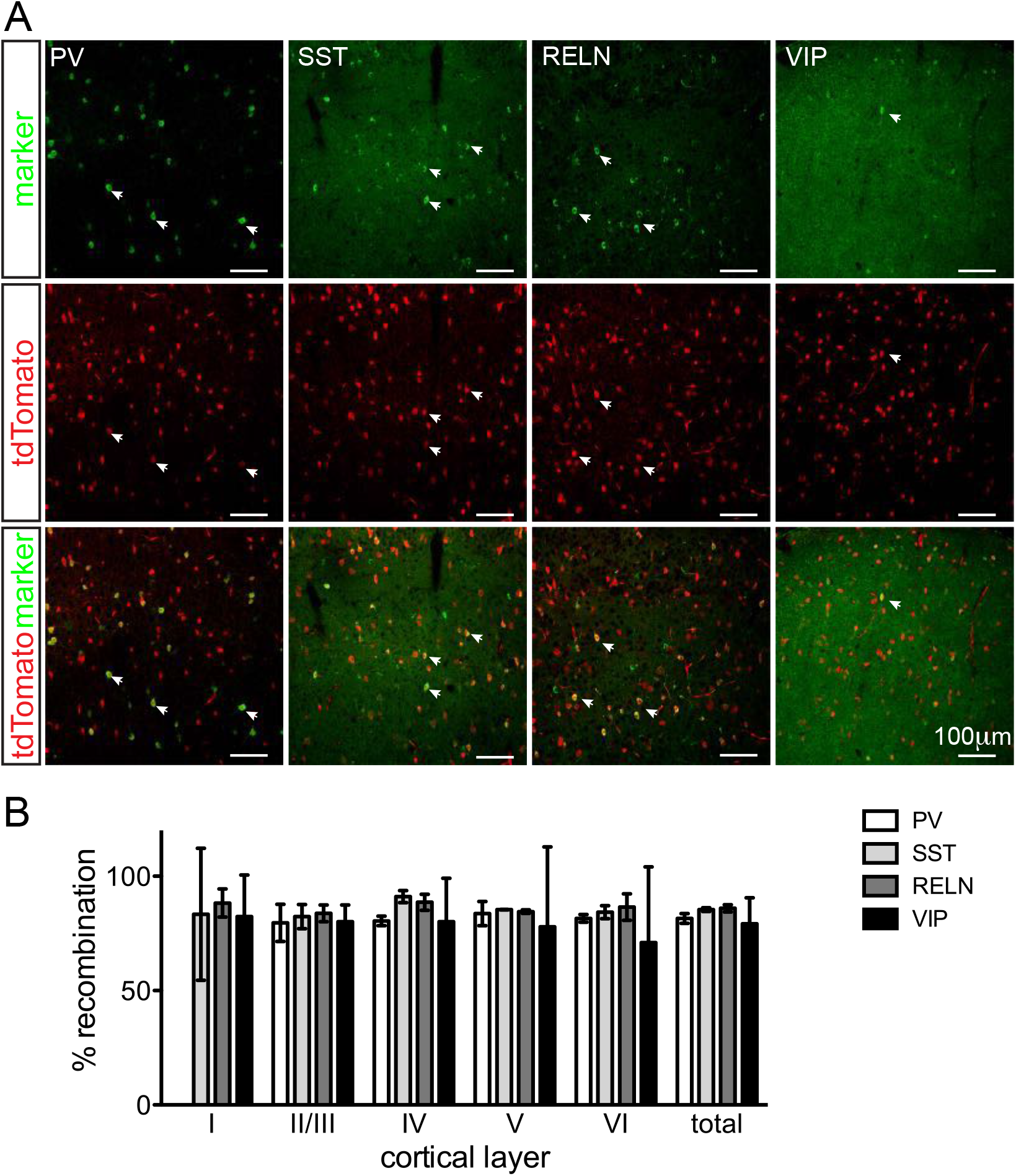
Assessment of recombination efficiency driven by Gad67Cre in cortical GABAergic interneurons. (A) Representative confocal images of fluorescence immunohistochemical detection of GABAergic interneuron markers (green) as indicated and tdTomato (red) in the somatosensory cortex of P30 Gad67Cre:Rosa26tdTom mice. Arrowheads point to a selection of cells co-expressing tdTomato and the respective marker. Scale bars: 100 µm. (B) Quantification of tdTomato+ cells co-expressing the indicated markers relative to the total of tdTomato+ cells counted. Results are expressed as average percentage ± SD. N=3 mice, 3 non-consecutive sections counted per mouse.

**Figure S5.**
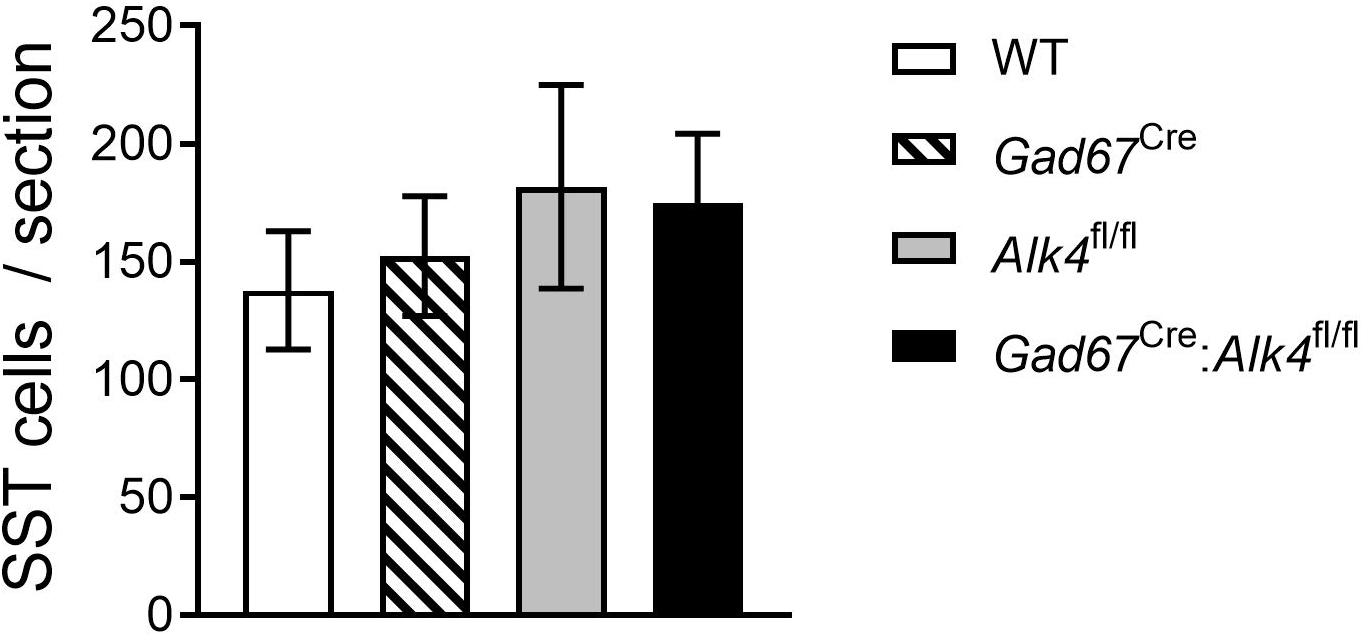
Assessment of SST+ cells in the striatum. Immunohistochemically labelled SST cells were counted in the striatum of P30 wild type (WT), Gad67Cre, Alk4fl/fl and Gad67Cre:Alk4fl/fl mice. Statistical analysis of the cell counts using ANOVA showed there was no difference in the number of SST cells between genotypes. 40 µm vibratome sections were collected between Bregma 1.7 --0.94 and every 6th section was fluorescence immunohistochemically labelled for SST. Cells were counted from pictures that were assembled from individual confocal images using the photomerge function in Adobe Photoshop.

**Figure S6.**
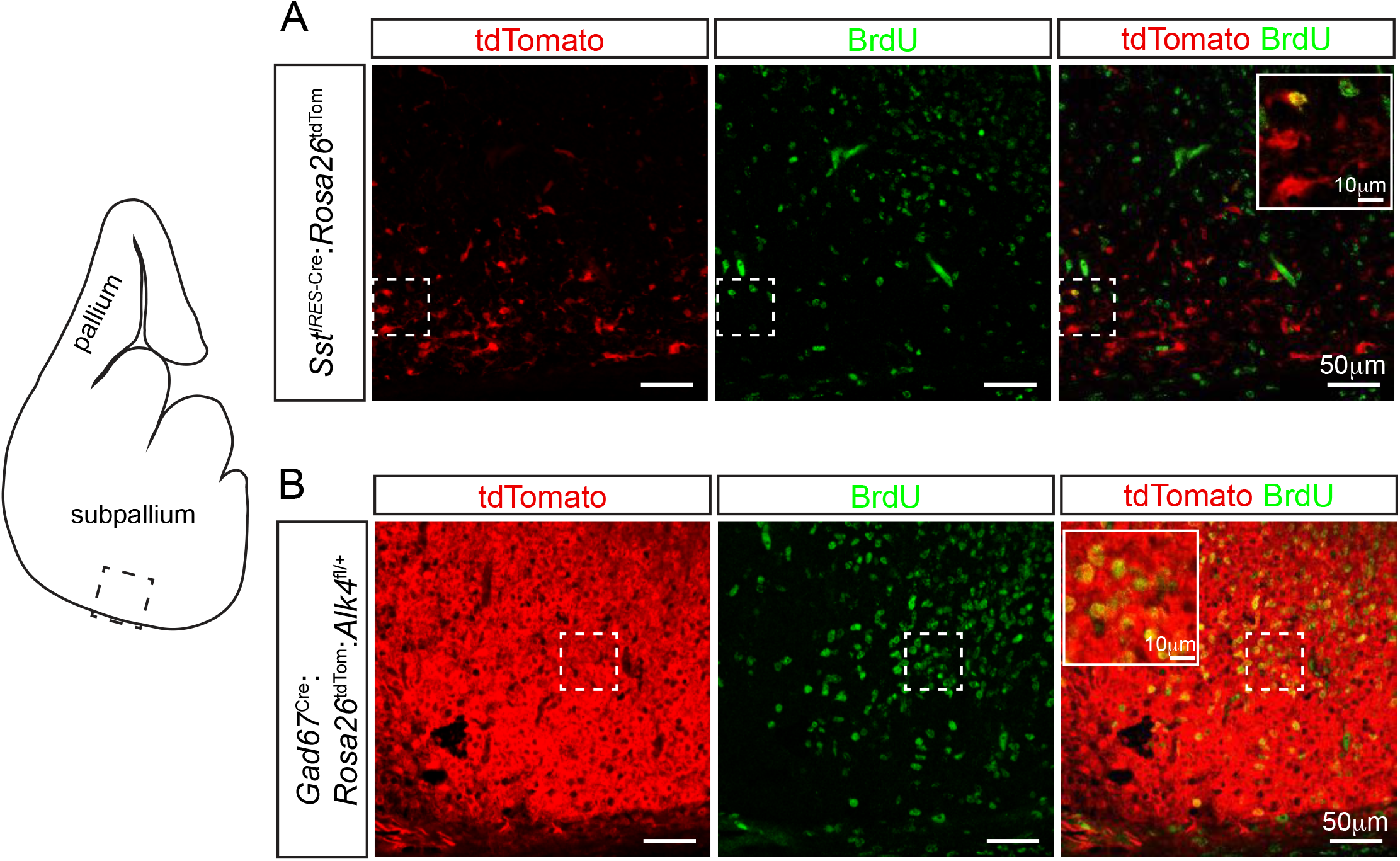
Comparison of tdTomato expression in SstIRES-Cre:Rosa26tdTom and Gad67Cre:Rosa26tdTom mice 48hs after BrdU labeling. Representative confocal images showing immunohistochemical detection tdTomato expression (red) driven by SstIRES-Cre (A) or Gad67Cre 48hs after BrdU labeling (green) in the subpallium (diagram) of E14.5 mouse embryos. Insets show a magnified view of the area outlined by a dashed box. Scale bars: 50 µm (main images), 10 µm (insets).

**Figure S7.**
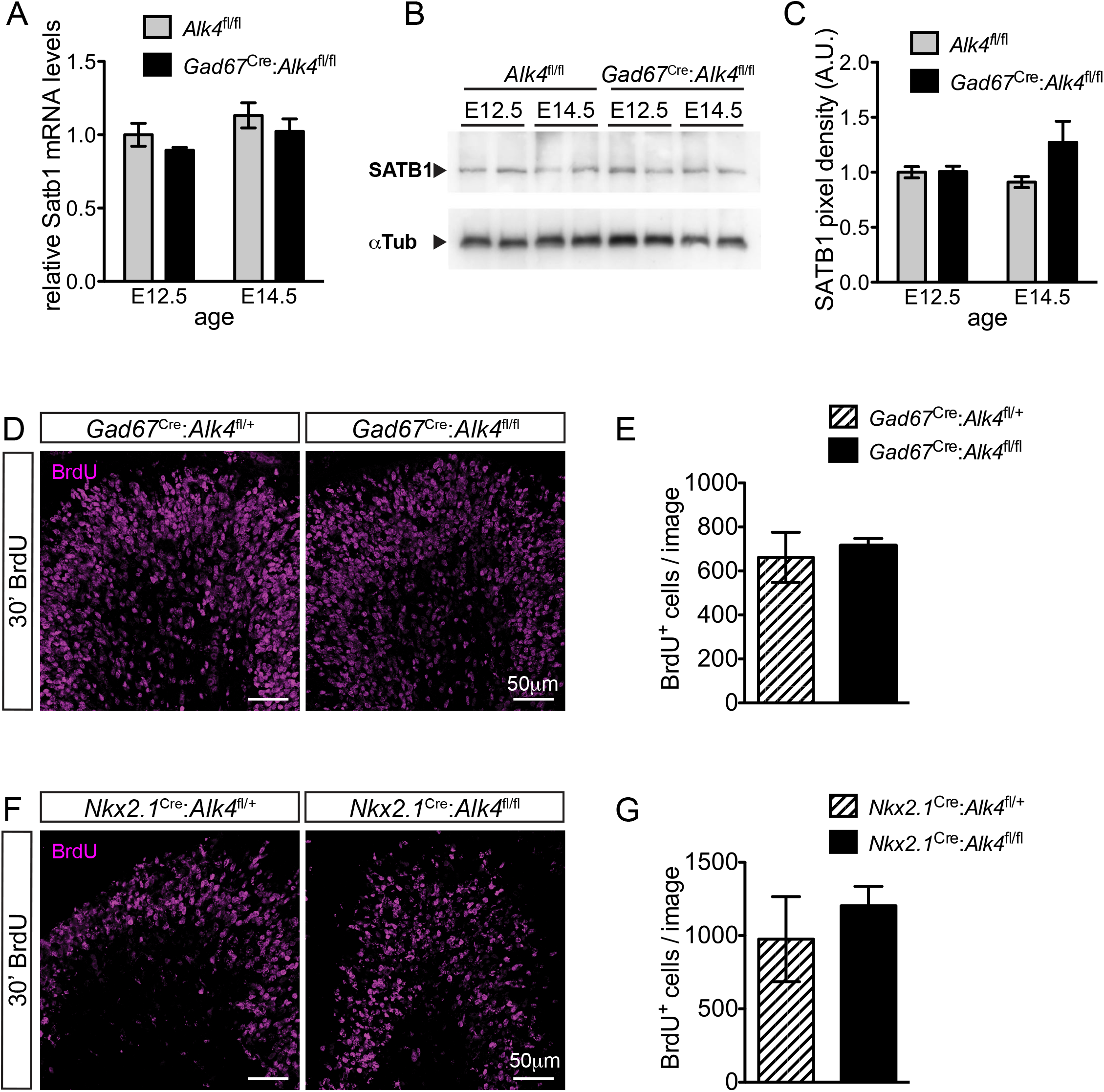
Unaltered Satb1 mRNA and protein levels and cell proliferation in the MGE of Alk4 mutant embryos. (A) Quantification of Satb1 mRNA expression by qRT-PCR analysis of E12.5 and E14.5 basal forebrain samples of Alk4fl/fl control (grey bars) and Gad67Cre:Alk4fl/fl (black bars) embryos. Results are presented as average ± SD. N=8 (E12.5 Alk4fl/fl), 6 (E12.5 Gad67Cre:Alk4fl/fl), 8 (E14.5 Alk4fl/fl), 7 (E14.5 Gad67Cre:Alk4fl/fl). There were no significant differences (ANOVA, Bonferroni post-hoc test). (B) Representative Western blot of whole cell lysates from E12.5 and E14.5 basal forebrain samples from Alk4fl/fl control and Gad67Cre:Alk4fl/fl embryos probed with antibodies to SATB1 (top) and αTubulin (bottom) as control. (C) Quantification of SATB1 protein expression relative to αTubulin in E12.5 and E14.5 basal forebrain samples from Alk4fl/fl control (grey bars) and Gad67Cre:Alk4fl/fl (black bars) embryos. Results are presented as average ± SD. N=4 embryos per genotype. There were no significant differences (ANOVA, Bonferroni post-hoc test). (D) Representative confocal images of immunohistochemical detection of BrdU (magenta) in MGE sections from E12.5 control (Gad67Cre:Alk4fl/+) and mutant (Gad67Cre:Alk4fl/fl) embryos. Embryos were harvested 30 min after the BrdU pulse. Scale bars: 50 µm. (E) Quantification of BrdU+ cells in the MGE of E12.5 control (Gad67Cre:Alk4fl/+) and mutant (Gad67Cre:Alk4fl/fl) embryos as indicated. Cell counts were normalized to the area of the section that was imaged and averaged per mouse. Results are presented as average ± SD. N=4 (Gad67Cre:Alk4fl/+), 7 (Gad67Cre:Alk4fl/fl). There were no significant differences (Student’s t-test). (F and G) Similar analysis to (E and F) on Nkx2.1Cre:Alk4fl/fl mutant embryos and controls. N=4 (Nkx2.1Cre:Alk4fl/+), 5 (Nkx2.1Cre:Alk4fl/fl). There were no significant differences (Student’s t-test).

**Figure S8.**
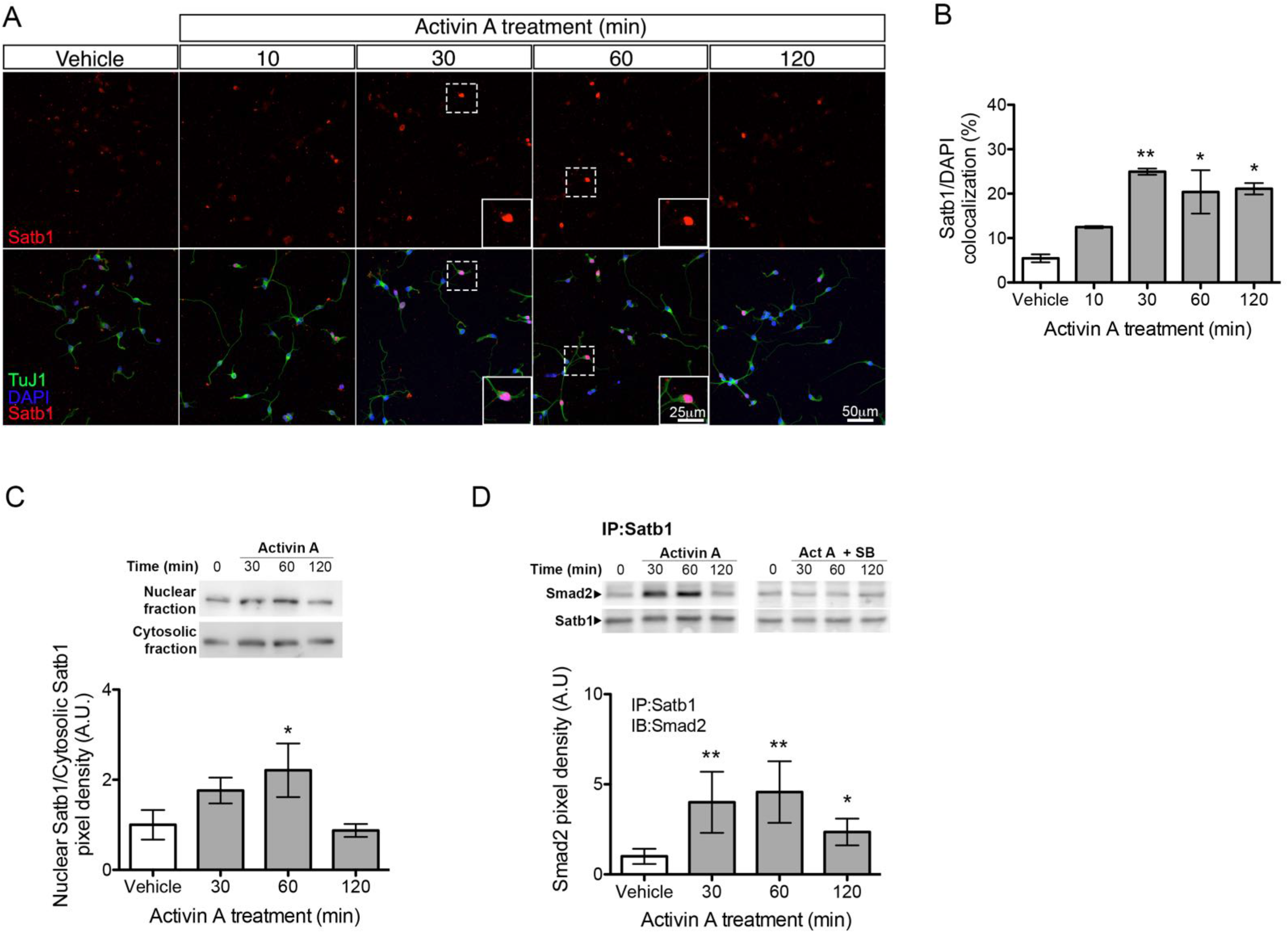
Activin A induces SATB1 nuclear localization and interaction with Smad 2 in MGE and Jurkat cell. (A) Representative confocal images of fluorescence immunohistochemical detection of SATB1 (green) in cultured MGE cells from wild type E14.5 embryos, counterstained for the neuronal marker Tuj1 (red) and DAPI (blue), after treatment with activin A for the indicated periods of time. Insets show magnified views of areas outlined by dashed boxes. Scale bars: 50 µm (main images), 25 µm (insets). (B) Quantification of E14.5 MGE cells displaying nuclear SATB1+ after activin A treatment. Results are expressed as percentage of the total cells (average ± SEM). N=5; *, P < 0.05; **, P < 0.01 compared to vehicle (One-Way ANOVA, P= 0.00957, followed (C) Representative Western blots of SATB1 in cytosolic and nuclear fractions prepared from Jurkat cells treated with Activin A for the indicated periods of time. Below, quantification of SATB1 in nuclear fractions relative to cytosolic fractions. Results are expressed as average ± SEM. N=4; *, P < 0.05 compared to vehicle (One-way ANOVA P=0.0235) followed, by (D) Representative Western blots of Smad2 and SATB1 in SATB1 immunoprecipitates prepared from Jurkat cells treated with Activin A with or without SB431542 (SB) for the indicated periods of time. Below, quantification of Smad2 co-immunoprecipitated by SATB1 antibodies normalized to SATB1. Results are expressed as average ± SEM. N=3; *, P < 0.05; **, P<0.01 compared to vehicle (One-way ANOVA P= 0.0042) followed, by

**Table S1.** P values.

| **Figure** | **Statistical test** | **P value** |
| --- | --- | --- |
| Figure 2B | Student's t-test<br>comparing the two<br>genotypes per layer | L1: P = 0.8969<br>L2/3: P = 0.0143<br>L4: P < 0.0001<br>L5: P = 0.0011<br>L6: P = 0.008<br>Total: P = 0.0007 |
| Figure 2D | ANOVA, Bonferroni<br>post-hoc test | L1: P = 0.0035<br>L2/3: P = 0.0972<br>L4: P = 0.006<br>L5: P = 0.823<br>L6: P = 0.9532<br>Total: P = 0.0088 |
| Figure 2E | ANOVA, Bonferroni<br>post-hoc test | L1: P = 0.9228<br>L2/3: P = 0.0012<br>L4: P = 0.0023<br>L5: P < 0.0001<br>L6: P < 0.0001<br>Total: P < 0.0001 |
| Figure 2F | ANOVA, Bonferroni<br>post-hoc test | L1: P = 0.1811<br>L2/3: P = 0.0055<br>L4: P = 0.0005<br>L5: P = 0.0002<br>L6: P < 0.0001<br>Total: P < 0.0001 |
| Figure 2G | ANOVA, Bonferroni<br>post-hoc test | L1: P = 0.5007<br>L2/3: P = 0.6143<br>L4: P = 0.1496<br>L5: P = 0.0149<br>L6: P = 0.1867<br>Total: P = 0.3389 |
| Figure 3B | ANOVA, Bonferroni<br>post-hoc test | L1: P = 0.9906<br>L2/3: P = 0.1911<br>L4: P = 0.0053<br>L5: P = 0.0193<br>L6: P = 0.0002<br>Total: P < 0.0001 |
| Figure 3D | Unpaired Student's t-test | L1: P = 0.2102<br>L2/3: P = 0.2088<br>L4: P = 0.4428<br>L5: P = 0.2439<br>L6: P = 0.2394<br>Total: P = 0.2645 |
| Figure 4B | Unpaired Student's t-test | L1: P = 0.5588<br>L2/3: P = 0.0035<br>L4: P = 0.0003<br>L5: P = 0.0024<br>L6: P < 0.0001<br>Total: P < 0.0001 |
| Figure 4C | Unpaired Student's t-test | L1: P = 0.9323<br>L2/3: P = 0.6004<br>L4: P = 0.1825<br>L5: P = 0.5771<br>L6: P = 0.0246<br>Total: P = 0.1991 |
| Figure 4D | Unpaired Student's t-test | L1: P = 0.0288<br>L2/3: P = 0.0001<br>L4: P = 0.0181<br>L5: P = 0.0479<br>L6: P = 0.0193<br>Total: P = 0.0025 |
| Figure 4E | Unpaired Student's t-test | L1: P = 0.9841<br>L2/3: P = 0.1358<br>L4: P < 0.0001<br>L5 P < 0.0001<br>L6: P = 0.0027<br>Total: P < 0.0001 |
| Figure 4F | Unpaired Student's t-test | L1: P = 0.0194<br>L2/3: P = 0.8241<br>L4: P = 0.1828<br>L5 P = 0.0538<br>L6: P = 0.0037<br>Total: P = 0.0373 |
| Figure 4G | Unpaired Student's t-test | L1: P = 0.1348<br>L2/3: P = 0.8454<br>L4: P = 0.2182<br>L5 P = 0.0016<br>L6: P = 0.1006<br>Total: P = 0.3086 |
| Figure 4I | Unpaired Student's t-test | L1: no SST<sup>+</sup>Chrna2<sup>+</sup> cells in this layer<br>L2/3: P = 0.3559<br>L4: no SST<sup>+</sup>Chrna2<sup>+</sup> cells in this layer<br>L5 P = 0.0079<br>L6: P = 0.0989<br>Total: P = 0.0135 |
| Figure 4K | Unpaired Student's t-test | L1: P = 0.6107<br>L2/3: P = 0.3152<br>L4: P = 0.1341<br>L5 P = 0.2149<br>L6: P = 0.6327<br>Total: P = 0.1534 |
| Figure 4M | Unpaired Student's t-test | L1: P = 0.3559<br>L2/3: P = 0.55<br>L4: P = 0.0054<br>L5 P = 0.0085<br>L6: P = 0.0368<br>Total: P = 0.0045 |
| Figure 5D | Unpaired Kolmogorov-Smirnov test<br>Unpaired Student's t-test (Amplitude and frequency) | Cumulative distribution: P < 0.0001<br>Amplitude: P = 0.003<br>Frequency: P = 0.024 |
| Figure 6B | Unpaired Student's t-test | L1: P = 0.6117<br>L2 - 4: P = 0.1296<br>L5 P = 0.0003<br>L6: P = 0.003<br>Total: P = 0.0007 |
| Figure 6C | Unpaired Student's t-test | L1: P = 0.8637<br>L2 - 4: P = 0.0061<br>L5 P = 0.0385 |
|  |  | L6: P = 0.6962<br>Total: P = 0.0842 |
| Figure 6E | Unpaired Student's t-test | MZ: P = 0.0074<br>L2 - 4: P = 0.0001<br>L5 P = 0.0001<br>L6: P = 0.0072<br>Total: P = 0.0001 |
| Figure 6F | Unpaired Student's t-test | MZ: P = 0.0216<br>L2 - 4: P = 0.6943<br>L5 P = 0.1261<br>L6: P = 0.0019<br>Total: P = 0.1826 |
| Figure 6I | Unpaired Student's t-test | Subpallium: P = 0.8871<br>Pallium: P = 0.9809 |
| Figure 7B | One-Way ANOVA<br>Bonferroni | Overall p=0.0083<br>Vehicle vs. 10 min p=0.8180<br>Vehicle vs. 30 min p=0.0051<br>Vehicle vs. 60 min p=0.0111<br>Vehicle vs. 120 min p=0.9815 |
| Figure 7D | One-Way ANOVA<br>Bonferroni multiple comparisons test | Overall p<0.001<br>Vehicle vs. Activin A p<0.0001<br>Vehicle vs. SB43152 p<0.05<br>Vehicle vs. Act + SB p>0.05 |
| Figure 7F | Two-way ANOVA<br>Bonferroni multiple comparisons test | Treatment p=0.022<br>Genotype p=0.0305<br>Interaction p=0.0305<br>WT Vehicle vs. WT Activin A p<0.01<br>KO Vehicle vs. KO Activin A p>0.05 |
| Figure 8B | One-Way ANOVA<br>Bonferroni multiple comparisons test | Overall p=0.01376<br>Vehicle vs. 30 min p<0.05<br>Vehicle vs. 90 min p>0.05 |
| Figure 8D | One-Way ANOVA<br>Bonferroni multiple comparisons test | Overall p<0.001<br>Vehicle vs. Activin A p<0.0001<br>Vehicle vs. SB43152 p>0.05<br>Vehicle vs. Act + SB p>0.05 |
| Figure 8F | Two-way ANOVA<br>Bonferroni multiple comparisons test | Treatment p=0.0369<br>Genotype p=0.0241<br>Interaction p=0.0055<br>WT Vehicle vs. WT Activin A p<0.01<br>KO Vehicle vs KO Activin A p>0. |
| Figure 9A | Student's t-test | p=0.035 |
| Figure 9B | Student's t-test | p=0.0431 |
| Figure S3A | Two way ANOVA<br>Bonferroni post-hoc test | Overall: P < 0.001<br>E12.5: P > 0.05<br>E14.5: P ≤ 0.05 |
| Figure S3B | Two way ANOVA<br>Bonferroni post-hoc test | Overall: P < 0.001<br>E12.5: P > 0.05<br>E14.5: P > 0.05 |
| Figure S3C | Two way ANOVA<br>Bonferroni post-hoc test | Overall: P < 0.001<br>E12.5: P > 0.05<br>E14.5: P > 0.05 |
| Figure S7A | Two-way ANOVA<br>Bonferroni | Overall: P = 0.227 |
| Figure S7C | Two-way ANOVA | Overall: P = 0.182 |
|  | Bonferroni |  |
| Figure S7E | Student's t-test | P = 0.4192 |
| Figure S7G | Student's t-test | P = 0.2829 |
| Figure S8B | One-Way ANOVA<br>Bonferroni | Overall: p=0.0096<br>Vehicle vs. 10 min p=0.3195<br>Vehicle vs 30 min p=0.0097<br>Vehicle vs 60 min p=0.0293<br>Vehicle vs 120 min p=0.0247 |
| Figure S8C | One-Way ANOVA<br>Bonferroni | Overall: p=0.0235<br>Vehicle vs. 30min p=0.3425<br>Vehicle vs. 60min: p=0.0415<br>Vehicle vs. 120min p=0.9934 |
| Figure S8D | One-Way ANOVA<br>Bonferroni | Overall: p p=0.0042<br>Vehicle vs. 30min p=0.0055<br>Vehicle vs. 60min: p=0.0152<br>Vehicle vs. 120min P=0.0395 |

## References

Alvarez, J.D., D.H. Yasui, H. Niida, T. Joh, D.Y. Loh, and T. Kohwi-Shigematsu. 2000. The MAR-binding protein SATB1 orchestrates temporal and spatial expression of multiple genes during T-cell development. Genes Dev. 14:521–535.

Andersen, F.G., J. Jensen, R.S. Heller, H.V. Petersen, L.I. Larsson, O.D. Madsen, and P. Serup. 1999. Pax6 and Pdx1 form a functional complex on the rat somatostatin gene upstream enhancer. FEBS Lett. 445:315–320.

Batista-Brito, R., R. Machold, C. Klein, and G. Fishell. 2008. Gene Expression in Cortical Interneuron Precursors is Prescient of their Mature Function. Cereb Cortex. 18:2306–2317. doi:10.1093/cercor/bhm258.

Budi, E.H., D. Duan, and R. Derynck. 2017. Transforming Growth Factor-β Receptors and Smads: Regulatory Complexity and Functional Versatility. Trends in Cell Biology. 27:658–672. doi:10.1016/j.tcb.2017.04.005.

Butt, S.J.B., V.H. Sousa, M.V. Fuccillo, J. Hjerling-Leffler, G. Miyoshi, S. Kimura, and G. Fishell. 2008. The requirement of Nkx2-1 in the temporal specification of cortical interneuron subtypes. Neuron. 59:722–732. doi:10.1016/j.neuron.2008.07.031.

Cai, S., H.-J. Han, and T. Kohwi-Shigematsu. 2003. Tissue-specific nuclear architecture and gene expression regulated by SATB1. Nat Genet. 34:42–51. doi:10.1038/ng1146.

Caputi, A., S. Melzer, M. Michael, and H. Monyer. 2013. The long and short of GABAergic neurons. Curr Opin Neurobiol. 23:179–186. doi:10.1016/j.conb.2013.01.021.

Close, J., H. Xu, N. De Marco García, R. Batista-Brito, E. Rossignol, B. Rudy, and G. Fishell. 2012. Satb1 is an activity-modulated transcription factor required for the terminal differentiation and connectivity of medial ganglionic eminence-derived cortical interneurons. J Neurosci. 32:17690–17705. doi:10.1523/JNEUROSCI.3583-12.2012.

Corbin, J.G., and S.J.B. Butt. 2011. Developmental mechanisms for the generation of telencephalic interneurons. Dev Neurobiol. 71:710–732. doi:10.1002/dneu.20890.

Corbin, J.G., S. Nery, and G. Fishell. 2001. Telencephalic cells take a tangent: non-radial migration in the mammalian forebrain. Nat Neurosci. 4 Suppl:1177–1182. doi:10.1038/nn749.

de Belle, I., S. Cai, and T. Kohwi-Shigematsu. 1998. The Genomic Sequences Bound to Special AT-rich Sequence-binding Protein 1 (SATB1) In Vivo in Jurkat T Cells Are Tightly Associated with the Nuclear Matrix at the Bases of the Chromatin Loops. J Cell Biol. 141:335–348. doi:10.1083/jcb.141.2.335.

Denaxa, M., M. Kalaitzidou, A. Garefalaki, A. Achimastou, R. Lasrado, T. Maes, and V. Pachnis. 2012. Maturation-promoting activity of SATB1 in MGE-derived cortical interneurons. Cell Rep. 2:1351–1362. doi:10.1016/j.celrep.2012.10.003.

Dickinson, L.A., C.D. Dickinson, and T. Kohwi-Shigematsu. 1997. An atypical homeodomain in SATB1 promotes specific recognition of the key structural element in a matrix attachment region. J Biol Chem. 272:11463–11470.

Dijke, ten, P., H. Yamashita, H. Ichijo, P. Franzén, M. Laiho, K. Miyazono, and C.-H. Heldin. 1994. Characterization of type I receptors for transforming growth factor-beta and activin. Science. 264:101–104.

Du, T., Q. Xu, P.J. Ocbina, and S.A. Anderson. 2008. NKX2.1 specifies cortical interneuron fate by activating Lhx6. Development. 135:1559–1567. doi:10.1242/dev.015123.

Flames, N., J.E. Long, A.N. Garratt, T.M. Fischer, M. Gassmann, C. Birchmeier, C. Lai, J.L.R. Rubenstein, and O. Marín. 2004. Short- and long-range attraction of cortical GABAergic interneurons by neuregulin-1. Neuron. 44:251–261. doi:10.1016/j.neuron.2004.09.028.

Flames, N., R. Pla, D.M. Gelman, J.L.R. Rubenstein, L. Puelles, and O. Marín. 2007. Delineation of multiple subpallial progenitor domains by the combinatorial expression of transcriptional codes. J Neurosci. 27:9682–9695. doi:10.1523/JNEUROSCI.2750-07.2007.

Forloni, G., C. Hohmann, and J.T. Coyle. 1990. Developmental expression of somatostatin in mouse brain. I. Immunocytochemical studies. Brain Res Dev Brain Res. 53:6–25.

Fuccillo, M., M. Rallu, A.P. McMahon, and G. Fishell. 2004. Temporal requirement for hedgehog signaling in ventral telencephalic patterning. Development. 131:5031–5040. doi:10.1242/dev.01349.

Gu, Z., M. Nomura, B.B. Simpson, H. Lei, A. Feijen, A. van den Eijnden-van Raaij, P.K. Donahoe, and E. Li. 1998. The type I activin receptor ActRIB is required for egg cylinder organization and gastrulation in the mouse. Genes Dev. 12:844–857.

Han, H.-J., J. Russo, Y. Kohwi, and T. Kohwi-Shigematsu. 2008. SATB1 reprogrammes gene expression to promote breast tumour growth and metastasis. Nature. 452:187–193. doi:10.1038/nature06781.

Hilscher, M.M., R.N. Leão, S.J. Edwards, K.E. Leão, and K. Kullander. 2017. Chrna2-Martinotti Cells Synchronize Layer 5 Type A Pyramidal Cells via Rebound Excitation. PLoS Biol. 15:e2001392. doi:10.1371/journal.pbio.2001392.

Hu, J.S., D. Vogt, M. Sandberg, and J.L. Rubenstein. 2017. Cortical interneuron development: a tale of time and space. Development. 144:3867–3878. doi:10.1242/dev.132852.

Kubota, Y., R. Hattori, and Y. Yui. 1994. Three distinct subpopulations of GABAergic neurons in rat frontal agranular cortex. Brain Res. 649:159–173.

Kumar, P.P., P.K. Purbey, C.K. Sinha, D. Notani, A. Limaye, R.S. Jayani, and S. Galande. 2006. Phosphorylation of SATB1, a global gene regulator, acts as a molecular switch regulating its transcriptional activity in vivo. Mol Cell. 22:231–243. doi:10.1016/j.molcel.2006.03.010.

Kumar, P.P., P.K. Purbey, D.S. Ravi, D. Mitra, and S. Galande. 2005. Displacement of SATB1-bound histone deacetylase 1 corepressor by the human immunodeficiency virus type 1 transactivator induces expression of interleukin-2 and its receptor in T cells. Mol Cell Biol. 25:1620–1633. doi:10.1128/MCB.25.5.1620-1633.2005.

Lim, L., D. Mi, A. Llorca, and O. Marín. 2018. Development and Functional Diversification of Cortical Interneurons. Neuron. 100:294–313. doi:10.1016/j.neuron.2018.10.009.

Liodis, P., M. Denaxa, M. Grigoriou, C. Akufo-Addo, Y. Yanagawa, and V. Pachnis. 2007. Lhx6 activity is required for the normal migration and specification of cortical interneuron subtypes. J Neurosci. 27:3078–3089. doi:10.1523/JNEUROSCI.3055-06.2007.

Lopez-Bendito, G., J.A. Sánchez-Alcañiz, R. Pla, V. Borrell, E. Picó, M. Valdeolmillos, and O. Marín. 2008. Chemokine signaling controls intracortical migration and final distribution of GABAergic interneurons. J Neurosci. 28:1613–1624. doi:10.1523/JNEUROSCI.4651-07.2008.

Ma, Y., H. Hu, A.S. Berrebi, P.H. Mathers, and A. Agmon. 2006. Distinct subtypes of somatostatin-containing neocortical interneurons revealed in transgenic mice. J Neurosci. 26:5069–5082. doi:10.1523/JNEUROSCI.0661-06.2006.

Marín, O., and J.L. Rubenstein. 2001. A long, remarkable journey: tangential migration in the telencephalon. 2:780–790. doi:10.1038/35097509.

Marín, O., and J.L.R. Rubenstein. 2003. Cell migration in the forebrain. Annu Rev Neurosci. 26:441–483. doi:10.1146/annurev.neuro.26.041002.131058.

Markram, H., M. Toledo-Rodriguez, Y. Wang, A. Gupta, G. Silberberg, and C. Wu. 2004. Interneurons of the neocortical inhibitory system. 5:793–807. doi:10.1038/nrn1519.

Massagué, J. 2012. TGFβ signalling in context. Nat Rev Mol Cell Biol. 13:616–630. doi:10.1038/nrm3434.

Mayer, C., C. Hafemeister, R.C. Bandler, R. Machold, R. Batista-Brito, X. Jaglin, K. Allaway, A. Butler, G. Fishell, and R. Satija. 2018. Developmental diversification of cortical inhibitory interneurons. Nature. 555:457–462. doi:10.1038/nature25999.

Mi, D., Z. Li, L. Lim, M. Li, M. Moissidis, Y. Yang, T. Gao, T.X. Hu, T. Pratt, D.J. Price, N. Sestan, and O. Marín. 2018. Early emergence of cortical interneuron diversity in the mouse embryo. Science. 360:81–85. doi:10.1126/science.aar6821.

Miyoshi, G., J. Hjerling-Leffler, T. Karayannis, V.H. Sousa, S.J.B. Butt, J. Battiste, J.E. Johnson, R.P. Machold, and G. Fishell. 2010. Genetic fate mapping reveals that the caudal ganglionic eminence produces a large and diverse population of superficial cortical interneurons. J Neurosci. 30:1582–1594. doi:10.1523/JNEUROSCI.4515-09.2010.

Montminy, M.R., and L.M. Bilezikjian. 1987. Binding of a nuclear protein to the cyclic-AMP response element of the somatostatin gene. Nature. 328:175–178. doi:10.1038/328175a0.

Moulder, R., T. Lönnberg, L.L. Elo, J.-J. Filén, E. Rainio, G. Corthals, M. Oresic, T.A. Nyman, T. Aittokallio, and R. Lahesmaa. 2010. Quantitative proteomics analysis of the nuclear fraction of human CD4+ cells in the early phases of IL-4-induced Th2 differentiation. Mol Cell Proteomics. 9:1937–1953. doi:10.1074/mcp.M900483-MCP200.

Moustakas, A., and C.-H. Heldin. 2009. The regulation of TGFbeta signal transduction. Development. 136:3699–3714. doi:10.1242/dev.030338.

Muñoz, W., R. Tremblay, D. Levenstein, and B. Rudy. 2017. Layer-specific modulation of neocortical dendritic inhibition during active wakefulness. Science. 355:954–959. doi:10.1126/science.aag2599.

Muñoz-Manchado, A.B., C. Bengtsson Gonzales, A. Zeisel, H. Munguba, B. Bekkouche, N.G. Skene, P. Lönnerberg, J. Ryge, K.D. Harris, S. Linnarsson, and J. Hjerling-Leffler. 2018. Diversity of Interneurons in the Dorsal Striatum Revealed by Single-Cell RNA Sequencing and PatchSeq. Cell Rep. 24:2179–2190.e7. doi:10.1016/j.celrep.2018.07.053.

Naka, A., J. Veit, B. Shababo, R.K. Chance, D. Risso, D. Stafford, B. Snyder, A. Egladyous, D. Chu, S. Sridharan, L. Paninski, J. Ngai, and H. Adesnik. 2018. Complementary networks of cortical somatostatin interneurons enforce layer specific control. bioRxiv. 1–99. doi:10.1101/456574.

Nakayama, Y., I.S. Mian, T. Kohwi-Shigematsu, and T. Ogawa. 2005. A nuclear targeting determinant for SATB1, a genome organizer in the T cell lineage. Cell Cycle. 4:1099–1106.

Narboux-Nême, N., R. Goïame, M.-G. Mattéi, M. Cohen-Tannoudji, and M. Wassef. 2012. Integration of H-2Z1, a somatosensory cortex-expressed transgene, interferes with the expression of the Satb1 and Tbc1d5 flanking genes and affects the differentiation of a subset of cortical interneurons. J Neurosci. 32:7287–7300. doi:10.1523/JNEUROSCI.6068-11.2012.

Nóbrega-Pereira, S., N. Kessaris, T. Du, S. Kimura, S.A. Anderson, and O. Marín. 2008. Postmitotic Nkx2-1 controls the migration of telencephalic interneurons by direct repression of guidance receptors. Neuron. 59:733–745. doi:10.1016/j.neuron.2008.07.024.

Pesold, C., W.S. Liu, A. Guidotti, E. Costa, and H.J. Caruncho. 1999. Cortical bitufted, horizontal, and Martinotti cells preferentially express and secrete reelin into perineuronal nets, nonsynaptically modulating gene expression. Proc Natl Acad Sci U S A. 96:3217–3222. doi:10.1073/pnas.96.6.3217.

Petilla Interneuron Nomenclature Group, G.A. Ascoli, L. Alonso-Nanclares, S.A. Anderson, G. Barrionuevo, R. Benavides-Piccione, A. Burkhalter, G. Buzsáki, B. Cauli, J. Defelipe, A. Fairén, D. Feldmeyer, G. Fishell, Y. Fregnac, T.F. Freund, D. Gardner, E.P. Gardner, J.H. Goldberg, M. Helmstaedter, S. Hestrin, F. Karube, Z.F. Kisvárday, B. Lambolez, D.A. Lewis, O. Marín, H. Markram, A. Muñoz, A. Packer, C.C.H. Petersen, K.S. Rockland, J. Rossier, B. Rudy, P. Somogyi, J.F. Staiger, G. Tamas, A.M. Thomson, M. Toledo-Rodriguez, Y. Wang, D.C. West, and R. Yuste. 2008. Petilla terminology: nomenclature of features of GABAergic interneurons of the cerebral cortex. Nature Reviews Neuroscience. 9:557–568. doi:10.1038/nrn2402.

Pozas, E., and C.F. Ibáñez. 2005. GDNF and GFRalpha1 promote differentiation and tangential migration of cortical GABAergic neurons. Neuron. 45:701–713. doi:10.1016/j.neuron.2005.01.043.

Schmierer, B., and C.S. Hill. 2007. TGFbeta-SMAD signal transduction: molecular specificity and functional flexibility. Nat Rev Mol Cell Biol. 8:970–982. doi:10.1038/nrm2297.

Southwell, D.G., M.F. Paredes, R.P. Galvao, D.L. Jones, R.C. Froemke, J.Y. Sebe, C. Alfaro-Cervello, Y. Tang, J.M. Garcia-Verdugo, J.L. Rubenstein, S.C. Baraban, and A. Alvarez-Buylla. 2012. Intrinsically determined cell death of developing cortical interneurons. Nature. 491:109–113. doi:10.1038/nature11523.

Taniguchi, H., M. He, P. Wu, S. Kim, R. Paik, K. Sugino, D. Kvitsiani, D. Kvitsani, Y. Fu, J. Lu, Y. Lin, G. Miyoshi, Y. Shima, G. Fishell, S.B. Nelson, and Z.J. Huang. 2011. A resource of Cre driver lines for genetic targeting of GABAergic neurons in cerebral cortex. Neuron. 71:995–1013. doi:10.1016/j.neuron.2011.07.026.

Tasic, B., V. Menon, T.N. Nguyen, T.-K. Kim, T. Jarsky, Z. Yao, B. Levi, L.T. Gray, S.A. Sorensen, T. Dolbeare, D. Bertagnolli, J. Goldy, N. Shapovalova, S. Parry, C. Lee, K. Smith, A. Bernard, L. Madisen, S.M. Sunkin, M. Hawrylycz, C. Koch, and H. Zeng. 2016. Adult mouse cortical cell taxonomy revealed by single cell transcriptomics. Nat Neurosci. 19:335–346. doi:10.1038/nn.4216.

Tasic, B., Z. Yao, L.T. Graybuck, K.A. Smith, T.N. Nguyen, D. Bertagnolli, J. Goldy, E. Garren, M.N. Economo, S. Viswanathan, O. Penn, T. Bakken, V. Menon, J. Miller, O. Fong, K.E. Hirokawa, K. Lathia, C. Rimorin, M. Tieu, R. Larsen, T. Casper, E. Barkan, M. Kroll, S. Parry, N.V. Shapovalova, D. Hirschstein, J. Pendergraft, H.A. Sullivan, T.-K. Kim, A. Szafer, N. Dee, P. Groblewski, I. Wickersham, A. Cetin, J.A. Harris, B.P. Levi, S.M. Sunkin, L. Madisen, T.L. Daigle, L. Looger, A. Bernard, J. Phillips, E. Lein, M. Hawrylycz, K. Svoboda, A.R. Jones, C. Koch, and H. Zeng. 2018. Shared and distinct transcriptomic cell types across neocortical areas. Nature. 563:72–78. doi:10.1038/s41586-018-0654-5.

Tomioka, R., K. Okamoto, T. Furuta, F. Fujiyama, T. Iwasato, Y. Yanagawa, K. Obata, T. Kaneko, and N. Tamamaki. 2005. Demonstration of long-range GABAergic connections distributed throughout the mouse neocortex. Eur J Neurosci. 21:1587–1600. doi:10.1111/j.1460-9568.2005.03989.x.

Verschueren, K., N. Dewulf, M.J. Goumans, O. Lonnoy, A. Feijen, S. Grimsby, K. Vandi Spiegle, P. ten Dijke, A. Morén, P. Vanscheeuwijck, C.H. Heldin, K. Miyazono, C. Mummery, J. Van Den Eijnden-Van Raaij, and D. Huylebroeck. 1995. Expression of type I and type IB receptors for activin in midgestation mouse embryos suggests distinct functions in organogenesis. Mech Dev. 52:109–123.

Wamsley, B., and G. Fishell. 2017. Genetic and activity-dependent mechanisms underlying interneuron diversity. Nature Reviews Neuroscience. 18:299–309. doi:10.1038/nrn.2017.30.

Wichterle, H., D.H. Turnbull, S. Nery, G. Fishell, and A. Alvarez-Buylla. 2001. In utero fate mapping reveals distinct migratory pathways and fates of neurons born in the mammalian basal forebrain. Development. 128:3759–3771.

Wonders, C.P., and S.A. Anderson. 2006. The origin and specification of cortical interneurons. 7:687–696. doi:10.1038/nrn1954.

Xu, H., H.-Y. Jeong, R. Tremblay, and B. Rudy. 2013. Neocortical somatostatin-expressing GABAergic interneurons disinhibit the thalamorecipient layer 4. Neuron. 77:155–167. doi:10.1016/j.neuron.2012.11.004.

Xu, Q., C.P. Wonders, and S.A. Anderson. 2005. Sonic hedgehog maintains the identity of cortical interneuron progenitors in the ventral telencephalon. Development. 132:4987–4998. doi:10.1242/dev.02090.

Xu, Q., M. Tam, and S.A. Anderson. 2008. Fate mapping Nkx2.1-lineage cells in the mouse telencephalon. J Comp Neurol. 506:16–29. doi:10.1002/cne.21529.

Yavorska, I., and M. Wehr. 2016. Somatostatin-Expressing Inhibitory Interneurons in Cortical Circuits. Front Neural Circuits. 10:76. doi:10.3389/fncir.2016.00076.

Zechel, S., P. Zajac, P. Lönnerberg, C.F. Ibáñez, and S. Linnarsson. 2014. Topographical transcriptome mapping of the mouse medial ganglionic eminence by spatially-resolved RNA-seq. Genome Biol. 15:486. doi:10.1186/PREACCEPT-9143209091271531.

Zeisel, A., A.B. Muñoz-Manchado, S. Codeluppi, P. Lönnerberg, G. La Manno, A. Juréus, S. Marques, H. Munguba, L. He, C. Betsholtz, C. Rolny, G. Castelo-Branco, J. Hjerling-Leffler, and S. Linnarsson. 2015. Brain structure. Cell types in the mouse cortex and hippocampus revealed by single-cell RNA-seq. Science. 347:1138–1142. doi:10.1126/science.aaa1934.

